# Chronic nicotine reduces nigral dopaminergic activity and remodels pedunculopontine cholinergic subpopulations

**DOI:** 10.64898/2026.02.12.705607

**Authors:** Rita Yu-Tzu Chen, Hassan Hosseini, Skylar Hall, Shirley Zhang, Rebekah Evans

**Author notes:** The authors declare no conflicts of interest.

## Abstract

Chronic nicotine is linked to neuroprotective effects and reduced Parkinson’s disease (PD) risk, yet its physiological effects on vulnerable substantia nigra pars compacta (SNc) dopaminergic and pedunculopontine nucleus (PPN) cholinergic neurons are not fully understood. These populations exhibit specialized properties such as spontaneous pacemaking and elevated dendritic calcium influx that contribute to their selective neurodegeneration in PD. Here, we used whole-cell patch-clamp, two-photon calcium imaging, and morphological reconstruction in acute brain slices from adult mice of both sexes following 8-10 weeks of oral nicotine. Chronic nicotine reduced spontaneous pacemaking, burst propensity, and rebound firing in SNc dopaminergic neurons, while decoupling tonic firing rate from proximal dendritic calcium levels. The changes in the SNc occurred without altering L-type channel contributions, dendritic arborization, or excitatory synaptic drive. In PPN cholinergic neurons, chronic nicotine induced subregion-specific plasticity. Rostral PPN neurons showed depolarized membrane potentials, broadened action potentials, and dendritic pruning, whereas caudal PPN neurons showed reduced spike frequency adaptation and accelerated excitatory postsynaptic current kinetics. These findings reveal potential cellular and circuit mechanisms by which nicotine may contribute to resilience against PD.

**Significant Statement:** Epidemiological studies have revealed nicotine’s surprising protective effect in Parkinson’s disease (PD). Interestingly, this protection appears only preventative, as nicotine shows no therapeutic benefits after PD diagnosis. Here, we evaluated the effects of chronic nicotine on two brainstem populations known to selectively degenerate in PD: substantia nigra pars compacta (SNc) dopaminergic neurons and pedunculopontine (PPN) cholinergic neurons. Chronic nicotine decreases activity levels of SNc neurons, depolarizes rostral PPN neurons, and enhances sustained firing capability in caudal PPN neurons. Notably, this is the first study to assess whether chronic nicotine alters the physiology and dendritic morphology of these vulnerable subpopulations. These cellular insights uncover potential mechanisms for promoting neuronal resilience in neurodegenerative diseases.

## Introduction

Nicotine is a toxic and addictive substance but has been linked to reduced risk of Parkinson’s disease (PD), neuroprotection, and improvement of cognitive symptoms in psychiatric disorders (Hernán et al., 2002; Kumari and Postma, 2005; Quik et al., 2008). In human epidemiology, nicotine use reduces the risk of developing PD by 30-70% (Hernán et al., 2002; Ritz et al., 2007; Breckenridge et al., 2016).

These protective effects appear to be only preventative, as nicotine does not reverse disease progression after neurodegeneration in patients or animal models (Huang et al., 2009; Thiriez et al., 2011; Oertel et al., 2023). Nicotine protection is dose-dependent, long-lasting (Hernán et al., 2002; Li et al., 2015; Breckenridge et al., 2016), dependent on nicotinic receptor activation, and requires intracellular calcium signaling (Stevens et al., 2003; Toulorge et al., 2011). These studies suggest that chronic nicotine induces long-term neuroplastic adaptations that reduce PD vulnerability. However, the detailed physiological mechanisms of nicotine’s beneficial actions remain unknown.

The degeneration of substantia nigra pars compacta (SNc) dopaminergic (DA) neurons is a hallmark of PD and causes extensive motor symptoms (Poewe et al., 2017; Gonzalez-Rodriguez et al., 2020). It has been hypothesized that several properties of SNc neurons make them particularly vulnerable: autonomous pacemaking, high levels of activity-dependent Ca^2+^ influx, extensive axonal arbor, and susceptibility to mitochondrial oxidative stress and α-synuclein aggregate accumulation (Giguère et al., 2018; Gonzalez-Rodriguez et al., 2020). In SNc DA neurons, L-type calcium channel-mediated Ca^2+^ entry during tonic firing (Chan et al., 2007; Sun et al., 2017) and T-type calcium channel-mediated influx during burst and rebound firing (Evans et al., 2017) have been implicated in cell-type specific vulnerability. Previous work in young mice found that chronic nicotine increases inhibitory drive onto SNc neurons, reducing their pacemaking activity (Nashmi et al., 2007; Xiao et al., 2009). Since PD risk increases sharply with age (Poewe et al., 2017), it is important to understand to understand how chronic nicotine influences SNc spontaneous and rebound firing when administered in adult mice.

While much previous research has focused on the role of SNc neurons in motor dysfunction, pedunculopontine nucleus (PPN) cholinergic (ACh) neurons also degenerate in PD, showing ∼30-60% loss in Parkinsonian patients (Jellinger, 1988; Rinne et al., 2008; Giguère et al., 2018; Sébille et al., 2019). PPN degeneration is linked to PD symptoms unresponsive to DA replacement therapy, including gait deficits, postural instability, cognitive impairments, and sleep disturbances (Perez-Lloret and Barrantes, 2016; Chambers et al., 2019). Midbrain cholinergic-dopaminergic connectivity is topographically organized, with the PPN projecting to the SNc and a closely related cholinergic nucleus, the laterodorsal tegmental nucleus (LDT), innervating the ventral tegmental area (VTA) (Gould et al., 1989; Lavoie and Parent, 1994; Oakman et al., 1995; Dautan et al., 2016). LDT-VTA circuitry is involved in nicotine-induced reward sensitization (Campos et al., 2025) and aversion (Wolfman et al., 2018; Liu et al., 2022). PPN ACh activity participates in nicotine reward prediction (Laviolette et al., 2002). Prenatal nicotine exposure alters the intrinsic properties of PPN neurons (Garcia-Rill et al., 2007) and acute nicotine hyperpolarizes PPN neurons in 21-day-old mice (Good et al., 2007). However, it is unknown how chronic nicotine modulates PPN ACh activity in adult animals. Understanding how adult-administered nicotine affects vulnerable neurons is critical for developing nicotine receptor- and circuit-based targets that could serve as preventative treatments for PD.

Here, we find that adult-administered chronic nicotine decreases overall activity in SNc DA neurons by dampening pacemaking frequency, rebound firing, and dendritic calcium-firing frequency relationship without altering the L-type calcium channel contribution. Chronic nicotine also induces subregion-specific plasticity in PPN ACh neurons. Specifically, rostral PPN neurons show depolarized membrane potential and reduced dendritic complexity, while caudal PPN neurons show reduced spike frequency adaptation and accelerated postsynaptic excitatory current kinetics. These findings reveal that chronic nicotine exerts cell type- and subregion-specific effects and may confer neuroprotection by remodeling the midbrain cholinergic-dopaminergic circuits.

## Materials and Methods

### Animals

All animal procedures were approved by the Georgetown University Medical Center Institutional Animal Care and Use Committee (IACUC). ChAT-Cre mice (Jax strain #031661) and Ai9-TdTomato mice (Jax strain #007909) on a C57BL/6J background were purchased from Jackson Laboratory and bred in the Georgetown University Department of Comparative Medicine (DCM) animal facility to produce heterozygous ChAT-Cre/Ai9-TdTomato offspring. The ChAT-Cre/Ai9 mice and wild-type (WT; Jax strain #000664) C57BL/6J mice were maintained under a 12 h light/12 h dark cycle with ad libitum access to food. Measures were taken to ensure minimal animal suffering and discomfort.

### Chronic nicotine administration

Nicotine administration was conducted across 20 independent cohorts of adult male and female mice (with treatment starting when mice were at least 8 weeks old), each beginning treatment on a separate start date. Most (18/20) cohorts consisted of 4 same-sex littermates, housed in pairs (2 assigned to control, 2 to nicotine). The remaining 2 cohorts included 3 control and 3 nicotine littermates, housed in trios; within each of these two cohorts, the control and nicotine mice were matched for sex and age (born within one week apart) to minimize variability related to developmental factors.

Nicotine was administered via home cage drinking water using custom-made water bottles as the sole water source. To habituate mice to the bitter taste of nicotine, we modified a previously established paradigm (Khwaja et al., 2007; Yang et al., 2023) by using 0.2% saccharin as vehicle and gradually increasing nicotine concentration. Nicotine (free base; Sigma-Aldrich) was dissolved in vehicle, which was protected from light and replaced every 3 days to prevent degradation. All mice initially received 0.2% saccharin for 6 days. The control group remained on the vehicle (saccharin). The nicotine group received escalating concentrations of nicotine (25, 50, 100, 200, and 300 µg/mL) at 3-day intervals, followed by maintenance at 300 µg/mL for 8–10 weeks. The final concentration was chosen because it produces maximum voluntary intake and plasma nicotine levels comparable to those observed in heavy smokers (Pekonen et al., 1993).

Fluid intake and body weight were measured every 3 days. Individual fluid intake was estimated as the average consumption per cage divided by the number of mice. Nicotine-containing water was continuously available until the time of electrophysiology or calcium imaging experiments to prevent withdrawal effects. Saccharin and nicotine treatments did not affect apparent health, as mice of both groups were active, showed normal nesting behavior, and maintained a shiny coat throughout the protocol.

### Electrophysiological solutions, brain slice preparation, and cell identification

Mice (average age: 211.0 ± 10.3 days; median age: 185 days) were anaesthetized with isoflurane and transcardially perfused with ice-cold slicing solution containing (in mM): 198 glycerol, 25 NaHCO_3_, 2.5 KCl, 1.2 NaH_2_PO_4_, 20 HEPES, 10 glucose, 10 MgCl_2_, 0.5 CaCl_2_ (bubbled with 95% O_2_/5% CO_2_, osmolarity = 310–320 mmol/kg). Horizontal brain slices (200 µm) were prepared using a Leica VT1200S vibratome in the slicing solution, and then were transferred to a tissue chamber and incubated for 30 mins at 34°C in holding solution containing (in mM): 92 NaCl, 30 NaHCO_3_, 2.5 KCl, 1.2 NaH_2_PO_4_, 20 HEPES, 35 glucose, 2 MgCl_2_, 2 CaCl_2_, 5 Na-ascorbate, 3 Na-pyruvate, 2 thiourea (bubbled with 95% O_2_/5% CO_2_, osmolarity = 300–310 mmol/kg). Slices were subsequently maintained at room temperature for at least 30 min before recording.

Whole-cell electrophysiology recordings were obtained from substantia nigra pars compacta (SNc) dopaminergic (DA) and pedunculopontine (PPN) cholinergic (ACh) neurons prepared from ChAT-Cre/Ai9 mice. Neurons were identified by anatomical landmarks and electrophysiological characteristics. The SNc was defined by its proximity to the substantia nigra pars reticulata (SNr) and its relative position to the medial terminal nucleus of the accessory optic tract (MT) (Masi et al., 2015; Yang et al., 2023). DA neurons were recognized by their large spindle-shaped somas located lateral to the MT, spontaneous pacemaking activity, and a prominent voltage sag in response to hyperpolarization (Grace and Onn, 1989; Masi et al., 2015).

The PPN was delineated by the distribution of ChAT^+^ cholinergic neurons (Rye et al., 1987), which exhibited tdTomato fluorescence and medium-to-large multipolar somas. Its boundaries were identified relative to the superior cerebellar peduncle (SCP) and laterodorsal tegmental nucleus (LDT). In ventral sections, the rostral PPN is located rostrolateral to the SCP, whereas the caudal PPN is mediocaudal to the SCP. In dorsal sections, the SCP clearly separates the laterally located caudal PPN from the medially located LDT.

Patch-clamp experiments were performed in a continuously perfused chamber at 30–34 °C using artificial cerebrospinal fluid (ACSF) containing (in mM): 125 NaCl, 25 NaHCO_3_, 3.5 KCl, 1.25 NaH_2_PO_4_, 10 glucose, 1 MgCl_2_ and 2 CaCl_2_ (bubbled with 95% O_2_/5% CO_2_, osmolarity = 295–310 mmol/kg). The internal pipette solution contained (in mM): 122 KMeSO_3_, 9 NaCl, 9 HEPES, 1.8 MgCl_2_, 14 phosphocreatine, 4 Mg-ATP, 0.3 Tris-GTP (pH = 7.35 with KOH; osmolarity = 290–300 mmol/kg), supplemented with 0.08-0.09% (weight/volume) neurobiotin for post-hoc labeling. Patch pipettes of tip resistance 2-6 MΩ were pulled from filamented borosilicate glass using a P-97 micropipette puller (Sutter Instrument). Recordings were made with a MultiClamp 700B amplifier and Digidata 1550B digitizer (Molecular Devices).

### Current-clamp recordings

Spontaneous firing, action potential (AP) waveform, afterdepolarization (ADP), rebound firing, and spike frequency adaptation (SFA) were recorded in whole-cell current-clamp configuration (sampling rate: 10 kHz, low-pass filter: 8 kHz). Bridge balance was adjusted, and liquid junction potential (–8 mV) was not corrected. Spontaneous pacemaking activity was recorded in gap-free mode without current injection and used for AP waveform analysis.

AP threshold potential was defined as 4% of the average maximum AP upstroke slope. The interspike membrane potential (Vm) was the mean Vm excluding spikes, measured between the AP threshold and full repolarization. The AP peak was identified as the amplitude of maximally depolarized Vm reached between upward crossing of an absolute voltage threshold (0 mV) and subsequent repolarization below a defined criterion (10 mV drop). The AP trough was defined as the most hyperpolarized potential after the AP peak. The AP half-width was defined as the time between 50% of the upstroke and 50% of the downstroke.

Rebound firing and SFA were measured using a series of eight current steps (–150 to +200 pA, 50 pA increments, 1 s duration). Rebound firing frequency was defined as the mean instantaneous firing frequency (inverse of interspike interval, ISI) within 500 ms after termination of the hyperpolarizing steps (–150 to –50 pA). The adaptation index was calculated as: 1 – (mean instantaneous frequency of the last 4 APs/mean instantaneous frequency of the first 4 APs) during the depolarizing steps (50–200 pA), where larger positive values indicate stronger adaptation (slowing of firing). Only steps with ³ 8 APs were included.

To record the hyperpolarization-induced ADP characteristic of ventral tier SNc neurons (Evans et al., 2017), SNc DA neurons were injected with hyperpolarizing current steps (2 s total duration) producing steady state potentials (SSP) ≤ –80 mV to allow deinactivation of T-type calcium channels. At the SSP, a transient depolarizing current step (1 nA, 5 ms) was used to elicit a single AP, followed immediately by an equal hyperpolarizing step. In ADP^+^ neurons, a large, slowly decaying depolarization would follow the AP before Vm returned to the SSP. ADP was quantified by measuring the area under the curve (AUC) of the ADP using trapezoidal integration.

### Voltage-clamp recordings

Spontaneous excitatory postsynaptic currents (sEPSCs) and M-type currents were measured in whole-cell voltage-clamp configuration (sampling rate: 100 kHz, low-pass filter: 6 kHz). Cell capacitance and serial resistance were 70% compensated and continuously monitored.

To record sEPSCs, neurons were voltage-clamped at –70 mV for 200 s. Using pClamp to analyze sEPSC parameters, 30–70 events were manually selected to create a detection template for each cell type and experimental condition (control and nicotine-treated SNc, rostral PPN, and caudal PPN). Events were then automatically detected using template matching (negative polarity, threshold = 4) and analyzed to obtain mean current amplitude, rise time, and decay time. Rise and decay time were defined as the time between the trace crossing 20% and 80% of the baseline-to-peak amplitude in the rising and falling stage, respectively. sEPSC frequency was calculated as the total number of detected events by recording duration.

M-currents were recorded using a protocol modified from previously described methods (Koyama and Appel, 2006; Bordas et al., 2015; Bayasgalan et al., 2021). Cells were depolarized to –20 mV to activate M-type potassium channels, followed by 1-s hyperpolarizing steps from –30 to –60 mV in 10 mV increments to induce deactivation. M-current amplitude was defined as the difference between the early and late steady state currents within the same voltage step, where contamination from other currents was minimal. Steps showing slow inward relaxations consistent with M-current deactivation were included, whereas steps exhibiting the opposite, outward relaxations were excluded from analysis. In total, 15 of 44 control PPN neurons and 16 of 51 nicotine PPN neurons had one or more steps excluded (number of steps excluded, breakdown by voltage: [control] –30 mV: 0; –40 mV: 4; –50 mV: 12; –60 mV: 16; [nicotine] –30 mV: 0; –40 mV: 7; –50 mV: 13; –60 mV: 18).

### Two-photon calcium imaging

Two-photon Ca^2+^ imaging of SNc DA neurons was performed on brain slices prepared from WT mice following previously published procedures (Evans et al., 2017; Chen and Evans, 2024). Cells were simultaneously patch-clamped in the recording chamber perfused with ACSF at 30–34 °C, as described above. After whole-cell break-in, cells were dialyzed for >15 min with the following internal solution (in mM): 122 KMeSO_3_, 9 NaCl, 9 HEPES, 1.8 MgCl_2_, 14 phosphocreatine, 4 Mg-ATP, 0.3 Tris-GTP, 0.05 Alexa Fluor 594, 0.3 Fluo-5F (pH = 7.35 with KOH; osmolarity = 290–300 mmol/kg). In cells that entered depolarization block (25 out of 47), a constant hyperpolarizing current (–5 to –160 pA; average = –67.2 pA) was applied throughout imaging to maintain regular pacemaking.

Neurons were imaged using a two-photon microscope (Bruker) equipped with a Mai Tai ultrafast Ti:sapphire laser (Spectra-Physics) tuned to 810 nm, which simultaneously excites Alexa Fluor 594 and Fluo-5F. Line scans (2 ms line period, 2 s duration) of somatodendritic regions were acquired through a 40×/0.8 NA objective (Olympus). Fluorescence was separated into red and green channels using standard dichroic and bandpass filters. Ca^2+^ signals were expressed as the green-to-red fluorescence ratio (G/R), normalized to the saturated ratio (Gs/R) obtained daily using pipettes containing the internal solution supplemented with 2 mM Ca^2+^.

Dendritic Ca^2+^ signals were measured at three sites in each neuron: soma, proximal dendrite (≤50 µm from soma), and distal dendrite (>50 µm from soma). Tonic Ca^2+^ signals were defined as the mean Ca^2+^ levels during slow tonic firing, whereas phasic Ca^2+^ signals were measured during burst-like firing evoked by a 200-pA current step and calculated as the peak minus baseline Ca^2+^ level in each trace. Ca^2+^ imaging data were quantified using ImageJ to determine fluorescence intensities and the distances of scanning sites from the soma. To inhibit L-type calcium channels, nifedipine (10 µM in DMSO; Tocris) was bath-applied for ³ 8 min before Ca^2+^ signal measurements.

### Immunohistochemistry

After electrophysiology and calcium imaging experiments, recorded brain slices were fixed in 4% (weight/volume) paraformaldehyde (PFA) in phosphate buffer (PB; pH = 7.6) for 12-24 hours, and then stored in PB at 4 °C. We used the hydrophilic tissue-clearing CUBIC method to increase tissue transparency (Susaki et al., 2014; Matsumoto et al., 2019) and combined it with our previously established immunofluorescence staining procedure (Evans et al., 2020; Fallah et al., 2025). All clearing steps were performed at room temperature on a shaker.

On day 1, slices were washed in PB (3 × 1 h) and incubated in CUBIC reagent 1 for 1–2 days. On day 3, slices were washed in PB (3 × 1 h), blocked in 0.5% fish gelatin solution for >3 h, and incubated in primary antibody in PB for 2 days. Slices containing the PPN were stained with streptavidin-Cy5 (Invitrogen SA1011; 1:1000) to label recorded neurons, whereas slices containing the SNc were co-stained with sheep anti-TH (Novus NB300-110; 1:1000) to label dopaminergic neurons and streptavidin-Cy5.

On day 5, slices were washed in PB (3 × 2 h) and incubated in secondary antibody in PB for 2 days. SNc slices were stained with donkey anti-sheep-568 (Invitrogen A11057; 1:1000) to visualize TH^+^ neurons with red fluorescence, while PPN slices remained in PB during this step. Finally, on day 7, slices were washed in PB (3 × 2 h), incubated in CUBIC reagent 2 for >2 h, and mounted on glass slides in reagent 2. Mounted tissues were sealed using adhesive frames (Thermo Scientific AB-0577) and glass coverslips, and stored in the dark before confocal imaging.

### Morphological reconstructions

Confocal imaging was performed on a Leica SP8 microscope equipped with a 20×/0.75 NA oil-immersion objective at Georgetown University Microscopy and Imaging Shared Resource core facility. Two fluorescence channels were collected sequentially: red (TH^+^ or ChAT^+^) and far red (neurobiotin-streptavidin). Image fields encompassing the entire soma and dendritic arbor of patched SNc and PPN neurons were acquired as tiled Z-stacks (1 µm step size, 1042 × 1042 pixels frame size, 400 Hz line scan rate). Z-stacks were compiled and exported as single TIFF files using ImageJ.

The TIFF files were converted and imported into Imaris (Bitplane) for three-dimensional reconstruction. For each cell, the soma was defined as the original starting point. The dendritic arbors were traced semi-automatically using the AutoPath algorithm with automatic center detection. To prevent double-tracing artifacts, each dendritic branch was initialized as a new path radiating outward from the soma. Sholl analysis was performed in Imaris with a radius step of 1 µm. Quantitative parameters including the number of intersections, total dendritic length, and soma volume were exported. The intersection counts per 10 µm radius were processed and graphed in Igor Pro. AUC of the Sholl curve was calculated using trapezoidal integration.

### Statistics

All numerical data are expressed as mean ± SEM. Capitalized N denotes the number of animals, and lowercase n denotes the number of cells per group. Igor Pro (WaveMetrics) was used for graphing and statistical analysis. Data normality was assessed using the Shapiro-Wilk test. Normally distributed data were analyzed using two-tailed t-test, and non-normal data using two-tailed Wilcoxon-Mann-Whitney (Wilcoxon) test; the smaller of the two W-statistic values is reported. Linear correlations were evaluated using Pearson’s correlation coefficient (r), and the difference between two correlation coefficients was tested using Fisher’s r-to-z transformation for independent correlations. Box plots display the median as the center line, 25^th^ and 75^th^ percentiles as the box, and 10^th^ and 90^th^ percentiles as the whiskers. For nicotine fluid intake data, Wilcoxon tests were performed with false discovery rate (FDR) correction; p-values were adjusted using the Benjamini-Hochberg method.

## Results

### Chronic oral nicotine temporarily reduces body weight and fluid intake

We established a chronic oral nicotine administration protocol in healthy adult mice by modifying an existing choice-free paradigm (Pekonen et al., 1993; Khwaja et al., 2007). Mice were pair-housed and given a single drinking water source containing 0.2% saccharin as the vehicle for 6 days as the baseline. In the nicotine treatment group, nicotine dose was started at 25 µg/mL and escalated every 3 days in the acclimation period until the maximum dose 300 µg/mL was reached and maintained for 8–10 weeks (Figure 1A).

**Figure 1.**
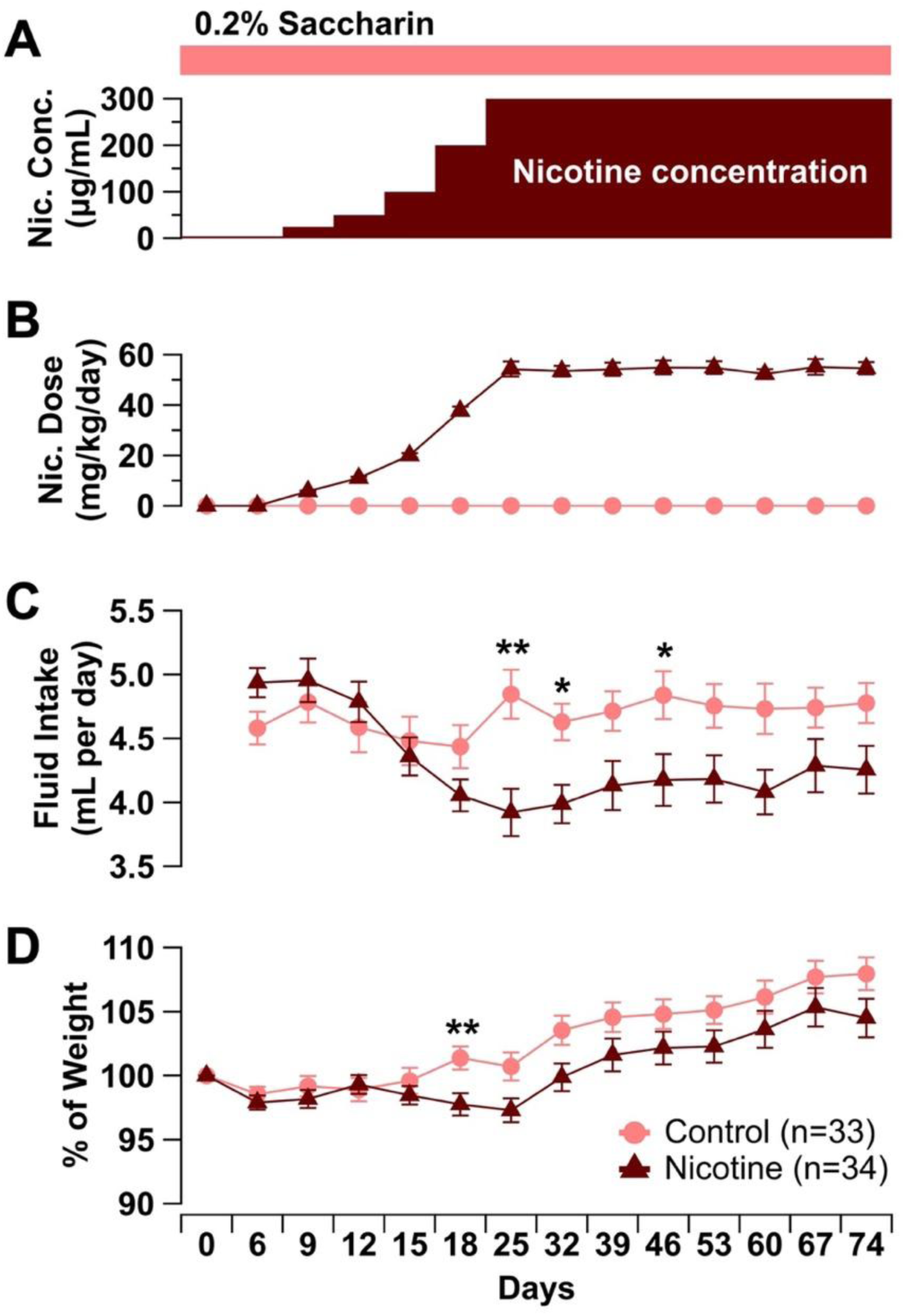
Chronic oral nicotine administration reduces fluid intake and body weight. **(A)** Nicotine dose escalation regimen: all mice initially received vehicle (0.2% saccharin) in drinking water for 6 days. One paired cage per cohort then was administered nicotine in vehicle starting at 25 μg/mL on day 6, increasing to 50 μg/mL on day 9, 100 μg/mL on day 12, 200 μg/mL on day 15, 300 μg/mL on day 18, and maintained at the final dose for 8-10 weeks (day 62-76). Control cages remained on vehicle throughout. **(B)** Estimated daily nicotine dose consumed per mouse, calculated as nicotine concentration in drinking water divided by daily fluid intake and body weight, plotted at 3-day intervals. **(C)** Estimated daily fluid intake per mouse, calculated from fluid consumed every three days divided by number of days and number of mice per cage. **(D)** Percent body weight change over time, normalized to day 0.

Mice in the control group (saccharin only) maintained stable fluid intake and age-associated weight increase. Nicotine treatment decreased fluid intake compared to control, reaching statistical significance at the 1^st^, 2^nd^, and 4^th^ week of 300 µg/mL (1^st^ week: W = 297, p = 0.0096; 2^nd^ week: W = 332, p = 0.0236; 4^th^ week: W = 355.5, p = 0.0407, control N = 33, nicotine N = 34, Wilcoxon test with FDR correction; Figure 1C). Despite the reduction of total fluid intake, nicotine intake (normalized to body weight) steadily increased over the escalation period and stabilized at 54.60 ± 2.42 mg/kg/day at the day of experiment (Figure 1B), a physiologically relevant dose comparable to that of heavy human smokers (Benowitz and Jacob, 1984).

Accompanying the decreased fluid intake, the body weight of nicotine-treated mice also decreased, though only reaching statistical significance at nicotine dose 200 µg/mL (W = 336.5, p = 0.0391, Wilcoxon test with FDR correction; Figure 1D). While the average fluid intake and body weight of mice in the nicotine group never completely recovered to the control level, the differences were not statistically significant during the chronic maintenance phase. This oral administration paradigm demonstrates that mice could be reliably maintained on a steady, clinically relevant dose of nicotine.

### Chronic nicotine reduces spontaneous and rebound firing frequency in SNc dopaminergic neurons

While nicotine may confer protection against PD through multiple mechanisms, one hypothesis is that chronic nicotine reduces the firing activity levels and excitability of SNc DA neurons, decreasing their overall metabolic load. To determine how chronic nicotine modifies these intrinsic electrophysiological properties of SNc DA neurons, we performed whole-cell patch-clamp recordings on acute brain slices from control and nicotine-treated mice. Nicotine-treated SNc DA neurons exhibited a significantly lower spontaneous pacemaking frequency than controls (control: 4.66 ± 0.32 Hz, N = 12, n = 36; nicotine: 3.70 ± 0.25 Hz, N = 12, n = 37; t = 2.37, p = 0.0204, t-test; Figure 2A,B), consistent with previous findings in young mice following a 10-day nicotine treatment paradigm (Nashmi et al., 2007; Xiao et al., 2009). We found that a subset of SNc DA neurons in both groups exhibited spontaneous burst firing in the absence of external stimulation. Interestingly, the proportion of bursting neurons was significantly lower in the nicotine group (3/38, 7.9%) than in controls (11/41, 26.8%; p = 0.0387, Fisher’s exact test; Figure 2C), in line with earlier reports that chronic nicotine decreases *in vivo* spontaneous burst firing in midbrain DA neurons in male rats (Grenhoff et al., 1991).

**Figure 2.**
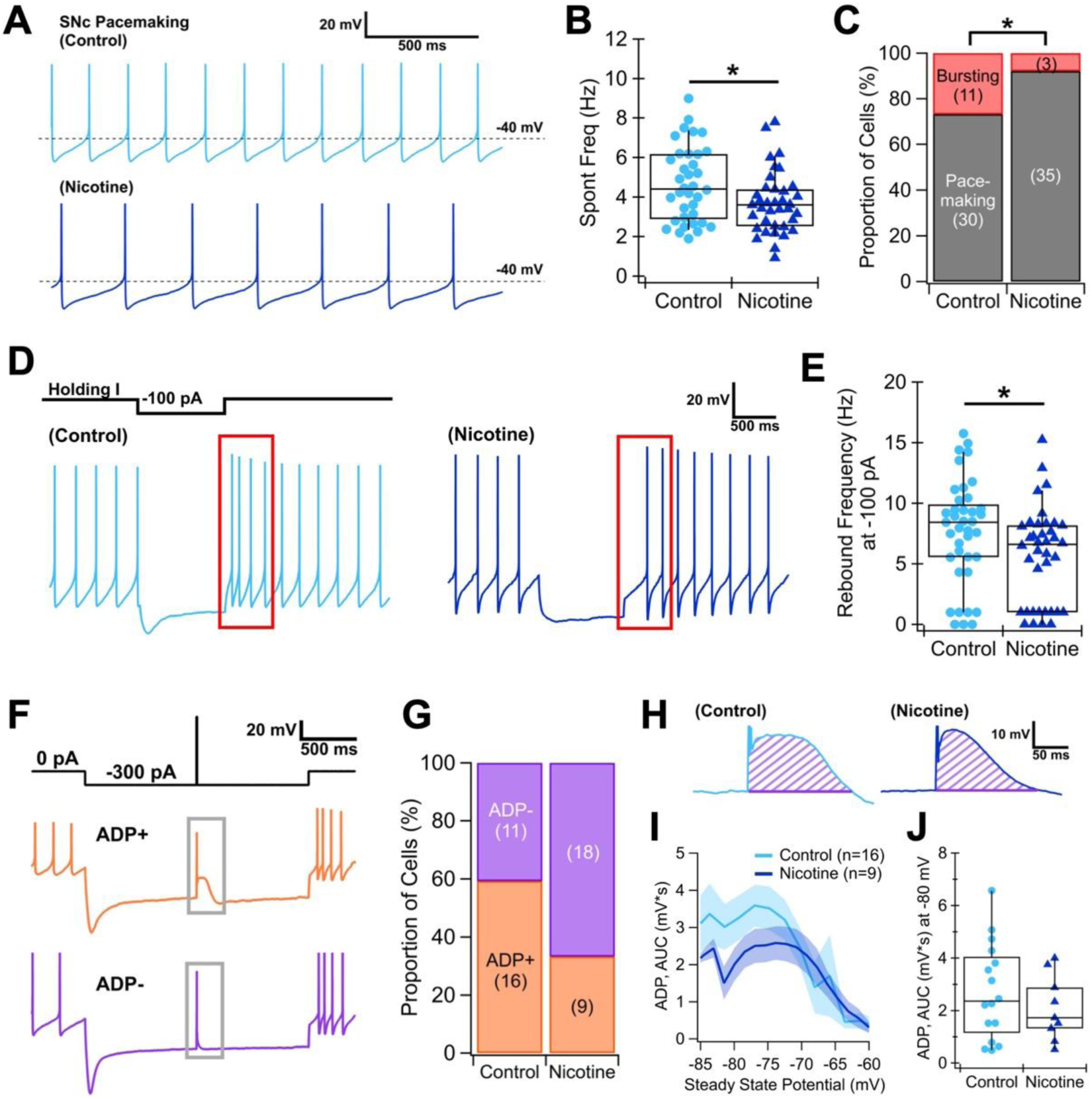
SNc dopaminergic (DA) neurons reduce excitability after chronic nicotine. **(A)** Representative traces showing spontaneous pacemaking of SNc DA neurons in control (light blue) and nicotine (dark blue) groups. **(B)** Summary of spontaneous pacemaking frequency. **(C)** Proportion of spontaneously bursting (red) and regularly pacemaking (gray) SNc DA neurons in control versus nicotine groups. **(D)** Representative rebound firing elicited by relief from a –100-pA hyperpolarizing current step. Action potentials within 500 ms post-hyperpolarization are labeled with red boxes and used for firing frequency analysis. **(E)** Summary of rebound firing frequency. **(F)** Representative hyperpolarization-induced afterdepolarization (ADP) recorded using a 2-s hyperpolarizing current step which produces a steady state potential (SSP) ≤ –80 mV, from which a transient depolarizing step is injected to elicit an action potential; an SNc DA neuron that shows prominent ADP response after action potential (ADP^+^, orange) and one where ADP is absent (ADP^-^, purple). **(G)** Proportion of ADP^+^ and ADP^-^ SNc DA neurons in control versus nicotine groups. **(H)** Representative ADP responses from ADP^+^ SNc DA neurons in control (light blue) and nicotine (dark blue) groups, with the area under the curve (AUC) of ADP shaded in purple stripes. **(I)** AUC of ADP plotted across the steady state potential reached during the 2-s hyperpolarizing step. **(J)** Summary of AUC of ADP at the steady state potential value closest to –80 mV.

Rebound activity is most prominent in the ventral tier SNc population, which is most vulnerable to neurodegeneration in PD (Yamada et al., 1990) and is accompanied by high levels of dendritic calcium influx (Evans et al., 2017). Therefore, rebound activity may be a source of SNc neuron vulnerability. Interestingly, nicotine-treated SNc DA neurons displayed a significantly lower rebound firing frequency following hyperpolarizing current injection (control 7.65 ± 0.69 Hz, n = 39; nicotine 5.61 ± 0.64 Hz, n = 38; W = 510, p = 0.0176, Wilcoxon test; Figure 2D,E). In the ventral tier SNc subpopulation, rebound firing is closely associated with a hyperpolarization-induced afterdepolarization (ADP), forming a distinct electrophysiological signature (Figure 2F) (Evans et al., 2017, 2020). To determine whether chronic nicotine alters this feature, we compared the proportion and average size of ADPs across groups. The fraction of ADP^+^ neurons was lower in the nicotine group (9/27, 33.3%) than in controls (16/27, 59.3%), though the difference was not statistically significant (p = 0.1007, Fisher’s exact test; Figure 2G).

Similarly, the average ADP magnitude trended smaller in nicotine-treated neurons but did not reach statistical significance (AUC of ADP at –80 mV, t = 1.02, p = 0.3193, t-test; Figure 2H–J). In contrast to spontaneous firing frequency and burst propensity, other firing properties in SNc DA neurons were not altered by nicotine exposure (Figure S1). Together, these results demonstrate that chronic nicotine dampens tonic, burst, and rebound activity in SNc DA neurons in adult mice of both sexes. Nicotine’s reduction of these three firing characteristics may decrease metabolic load of SNc neurons and thereby enhance resilience to mitochondrial stress and other pathological perturbations (such as α-synuclein aggregation) associated with Parkinsonian degeneration.

### Chronic nicotine decouples the relationship between tonic firing frequency and dendritic calcium signals in SNc dopaminergic neurons

Because elevated intracellular calcium contributes to neuronal vulnerability (Dryanovski et al., 2013; Sun et al., 2017; Guzman et al., 2018), we examined chronic nicotine effects on calcium dynamics in SNc DA neurons. Two-photon linescan imaging was performed at the soma, proximal dendrite, and distal dendrite during action potential activity (Figure 3A). Tonic calcium signals were measured during spontaneous pacemaking at 0 pA, whereas phasic calcium signals were recorded during a 200-pA depolarizing step that elicits burst-like firing, as in (Chen and Evans, 2024) (Figure 3B). Tonic calcium levels did not differ significantly between control and nicotine-treated neurons at the soma (control, N = 15, n = 25; nicotine, N = 14, n = 22; W = 256, p = 0.6960, Wilcoxon test; Figure 3C), proximal dendrite (control, n = 25; nicotine, n = 22; t = -0.58, p = 0.5669, t-test; Figure 3D), or distal dendrite (control, n = 24; nicotine, n = 22; t = -0.32, p = 0.7514, t-test; Figure 3E).

**Figure 3.**
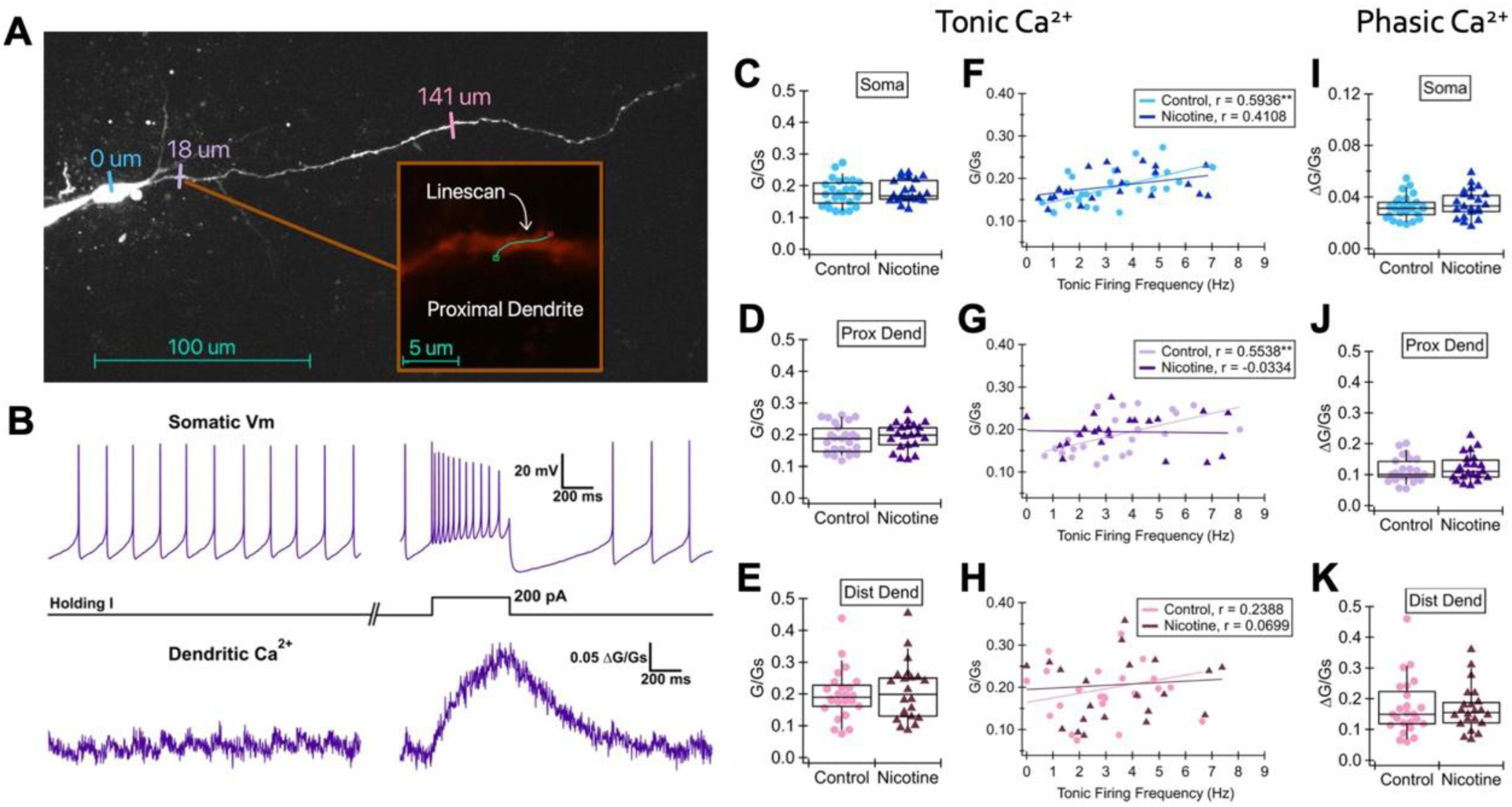
SNc DA neurons show no changes in tonic and phasic dendritic calcium signal amplitudes but decoupling of tonic calcium signals from firing rates after chronic nicotine. **(A)** Representative two-photon Z-stack of a SNc DA neuron with a patch pipette (left) filled with Alexa Fluor 594 and Fluo-5F and sites where linescans were taken: soma (0 μm), proximal dendrite (≤50 μm), and distal dendrite (>50 μm); an inset (orange box) from the area indicated by the orange line pointing away from the cell, showing the site of linescan taken at the proximal dendrite. **(B)** Representative time-matched whole-cell somatic Vm recording (top) and dendritic linescan Ca^2+^ signal (bottom) during 0 pA holding current (left) and a 200-pA current step (right) **(C)** Summary of the mean Ca^2+^ signals during tonic firing measured at the soma, **(D)** proximal dendrite, and **(E)** distal dendrite. **(F)** Mean tonic Ca^2+^ signals measured at the soma, **(G)** proximal dendrite, and **(H)** distal dendrite plotted against the tonic firing frequencies. Data from each treatment group at each location were fitted with linear regression lines. The corresponding Pearson correlation coefficients (r) are shown. **(I)** Summary of phasic Ca^2+^ signals (peak – baseline) during the 200-pA current step measured at the soma, **(J)** proximal dendrite, and **(K)** distal dendrite.

Importantly, this imaging method does not allow for calcium concentration measurements, thus the data presented here are relative to saturated values rather than absolute millimolar concentrations. However, previous work shows that tonic firing frequency correlates with dendritic calcium levels in SNc DA neurons (Hage and Khaliq, 2015). As expected, this positive correlation was evident in control neurons at the soma (r = 0.59, p = 0.0018; Figure 3F) and proximal dendrite (r = 0.55, p = 0.0041; Figure 3G). Strikingly, chronic nicotine flattened the slope of this relationship in the proximal dendrites (r = – 0.03, p = 0.8827), indicating reduced calcium influx in cells with higher firing frequencies (r_control vs r_nicotine, p = 0.0358, Fisher’s r-to-z test; Figure 3G). While a slight flattening of this relationship with chronic nicotine was also observed in the soma (Figure 3F) and distal dendrites (Figure 3H), the correlations were not significantly different from those of control. Phasic calcium transients did not differ between the control and nicotine group at the soma (p = 0.3890, t = -0.87, t-test; Figure 3I), proximal dendrite (p = 0.5754, W = 248, Wilcoxon test; Figure 3J), or distal dendrite (p = 0.8703, W = 256, Wilcoxon test; Figure 3K).

Because low-threshold L-type calcium channels are major contributors to tonic calcium influx in SNc DA neurons (Chan et al., 2007; Guzman et al., 2009; Hage and Khaliq, 2015), we tested whether their activity was altered by chronic nicotine. Bath application of the L-type calcium channel blocker nifedipine (10 µM), as in (Chen and Evans, 2024), revealed no significant nicotine-induced differences in either tonic or phasic calcium components (Figure S2). Together, these data indicate that chronic nicotine selectively weakens the coupling between tonic firing rate and calcium influx in proximal dendrites, but does not reduce total dendritic calcium signals and calcium influx through L-type channels.

### Chronic nicotine does not alter dendritic morphology or excitatory synaptic inputs onto SNc dopaminergic neurons

In addition to modulation of intrinsic excitability and calcium dynamics, nicotine could influence neuronal resilience through structural or synaptic remodeling. Chronic nicotine-induced morphological plasticity occurs in striatal projection neurons (McDonald et al., 2005; Ehlinger et al., 2016, 2017).

However, no previous studies have directly examined nicotine’s influence on dendritic morphology in SNc DA neurons, despite their central role in both nicotine addiction (Picciotto et al., 1998; Nashmi et al., 2007) and PD pathology (Gonzalez-Rodriguez et al., 2020). To determine whether chronic nicotine alters dendritic architecture, we performed Sholl analysis on neurobiotin-filled neurons reconstructed after whole-cell recording (Figure 4A,B). Total dendritic length did not differ significantly between control and nicotine-treated groups (control: N = 15, n = 32; nicotine: N = 15, n = 38; t = 0.24, p = 0.8114, t-test; Figure 4C). Likewise, the number of dendritic intersections as a function of distance from the soma showed fully overlapping profiles, indicating no effect of chronic nicotine on SNc dendritic complexity (Figure 4D).

**Figure 4.**
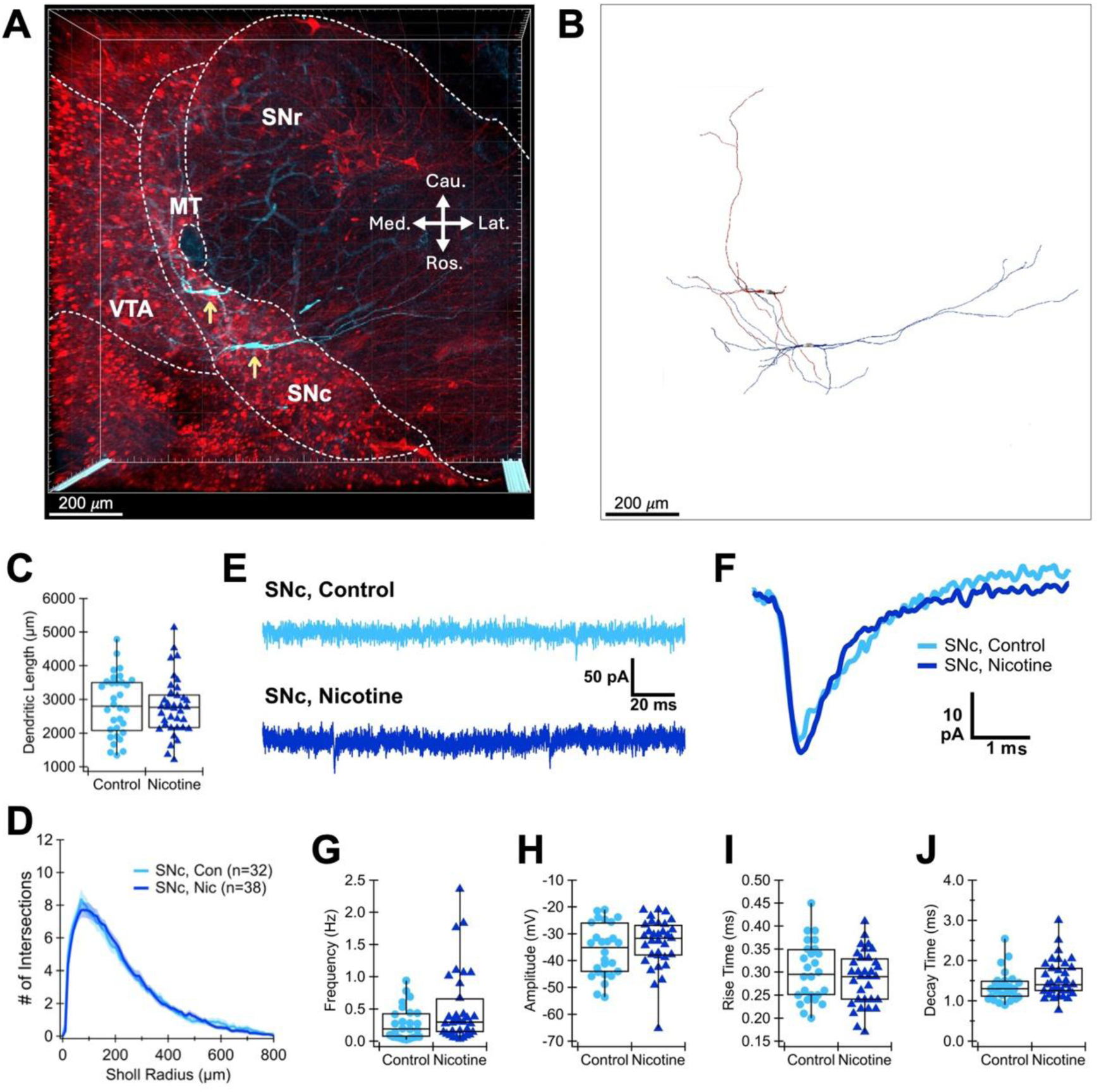
SNc DA neurons show no changes in spontaneous excitatory postsynaptic current (sEPSC) properties and dendritic morphology after chronic nicotine. **(A)** Three-dimensional confocal Z-stack image reconstructed in Imaris of two patched SNc DA neurons (cyan, neurobiotin-filled, streptavidin-Cy5 stain), with yellow arrows indicating somas, and neighboring DA neurons (red, TH^+^ antibody stain) on a horizontal brain slice; dashed lines delineate approximate boundaries of landmarks: SNr (substantia nigra pars reticulata), MT (medial terminal nucleus of the accessory optic tract), VTA (ventral tegmental area), and SNc (substantia nigra pars compacta). Cau = caudal, Med = medial, Ros = rostral, and Lat = lateral. **(B)** The same two SNc DA neurons shown in (A) with soma and dendrites digitally reconstructed in Imaris. **(C)** Summary of total dendritic length of SNc DA neurons in control (light blue) and nicotine (dark blue) groups. **(D)** Sholl analysis of SNc DA neurons, with the number of dendritic intersections plotted against Sholl radius value of 10 μm. **(E)** Representative voltage-clamp recording of sEPSCs at –70 mV holding potential. **(F)** Representative averaged sEPSC waveform from one control (light blue) and one nicotine (dark blue) SNc DA neurons. **(G)** Summary of sEPSC frequency, **(H)** amplitude, **(I)** rise time, **(J)** decay time.

Previous work has shown that chronic nicotine increases inhibitory synaptic inputs to SNc DA neurons (Nashmi et al., 2007; Xiao et al., 2009), but its effects on excitatory synaptic inputs remain unknown. In other brain regions, chronic nicotine alters glutamatergic signaling in models using younger animals. For example, it downregulates AMPA receptor GluA2/3 subunits in the striatum and VTA (Adriani et al., 2004; Vieyra-Reyes et al., 2008), while both acute and chronic nicotine exposure increases AMPA/NMDA ratio in VTA DA neurons (Gao et al., 2010; Campos et al., 2025). Given our findings that chronic nicotine decreases spontaneous firing frequency and burst propensity in adult SNc neurons, we hypothesized that excitatory synaptic drive might also be reduced. To test this, we recorded spontaneous excitatory postsynaptic currents (sEPSCs) from SNc DA neurons (Figure 4E,F). The average sEPSC frequency, amplitude, rise time, and decay time did not differ significantly between control and nicotine-treated groups (control: N = 11, n = 26; nicotine: N = 12, n = 33; frequency: W = 331.5, p = 0.1383, Wilcoxon test; amplitude: W = 348, p = 0.2209, Wilcoxon test; rise time: t = 0.86, p = 0.3925, t-test; decay time: W = 320.5, p = 0.0984, Wilcoxon test; Figure 4G–J).

Because the effects of nicotine might be confined to specific subpopulations of SNc neurons, we separately analyzed ADP^+^ and ADP^-^ SNc DA neurons. Sholl analysis and sEPSC properties of control and nicotine-treated neurons did not differ within either subtype (Figure S3). Together, these data indicate that chronic nicotine does not alter dendritic arborization or excitatory synaptic inputs to SNc DA neurons, even when accounting for cellular heterogeneity. The absence of structural or excitatory synaptic changes suggests that nicotine-induced reductions in spontaneous and rebound firing (Figure 2) and attenuation of the firing rate-calcium relationship (Figure 3) are not secondary to dendritic remodeling or due to loss of excitatory drive.

### Chronic nicotine selectively alters action potential shape in rostral PPN cholinergic neurons

Two mesopontine cholinergic nuclei, the pedunculopontine nucleus (PPN) and lateral dorsal tegmental nucleus (LDT), provide the principal cholinergic inputs to midbrain DA neuron cell bodies and dendrites in a topographically organized manner, with PPN projections targeting the SNc and LDT projections innervating the VTA (Martinez-Gonzalez et al., 2011; Mena-Segovia and Bolam, 2017; Xiao et al., 2020). The PPN is also a selective site of vulnerability in PD, showing around 40–60% of cholinergic cell loss (Giguère et al., 2018), highlighting the importance of characterizing how chronic nicotine influences PPN physiology. Chronic nicotine disrupts LDT cholinergic modulation of VTA DA neurons and alters motivational behaviors (Campos et al., 2025), but nicotine’s effects on the PPN-SNc pathway remain unknown. Therefore, we examined the effects of chronic nicotine on the electrophysiological properties of PPN ACh neurons. We compared spontaneous firing and action potential (AP) properties from control and nicotine-treated mice. The PPN can be subdivided into rostral and caudal regions (Figure 5A), which differ in electrophysiological characteristics, synaptic inputs, and projection patterns (Martinez-Gonzalez et al., 2011; Baksa et al., 2019; Fallah et al., 2025). Therefore, we analyzed the rostral and caudal PPN separately. Across both subregions, spontaneous firing frequency, firing regularity, input resistance, whole-cell capacitance, and current-frequency relationship did not differ significantly between control and nicotine groups (Figure S4).

**Figure 5.**
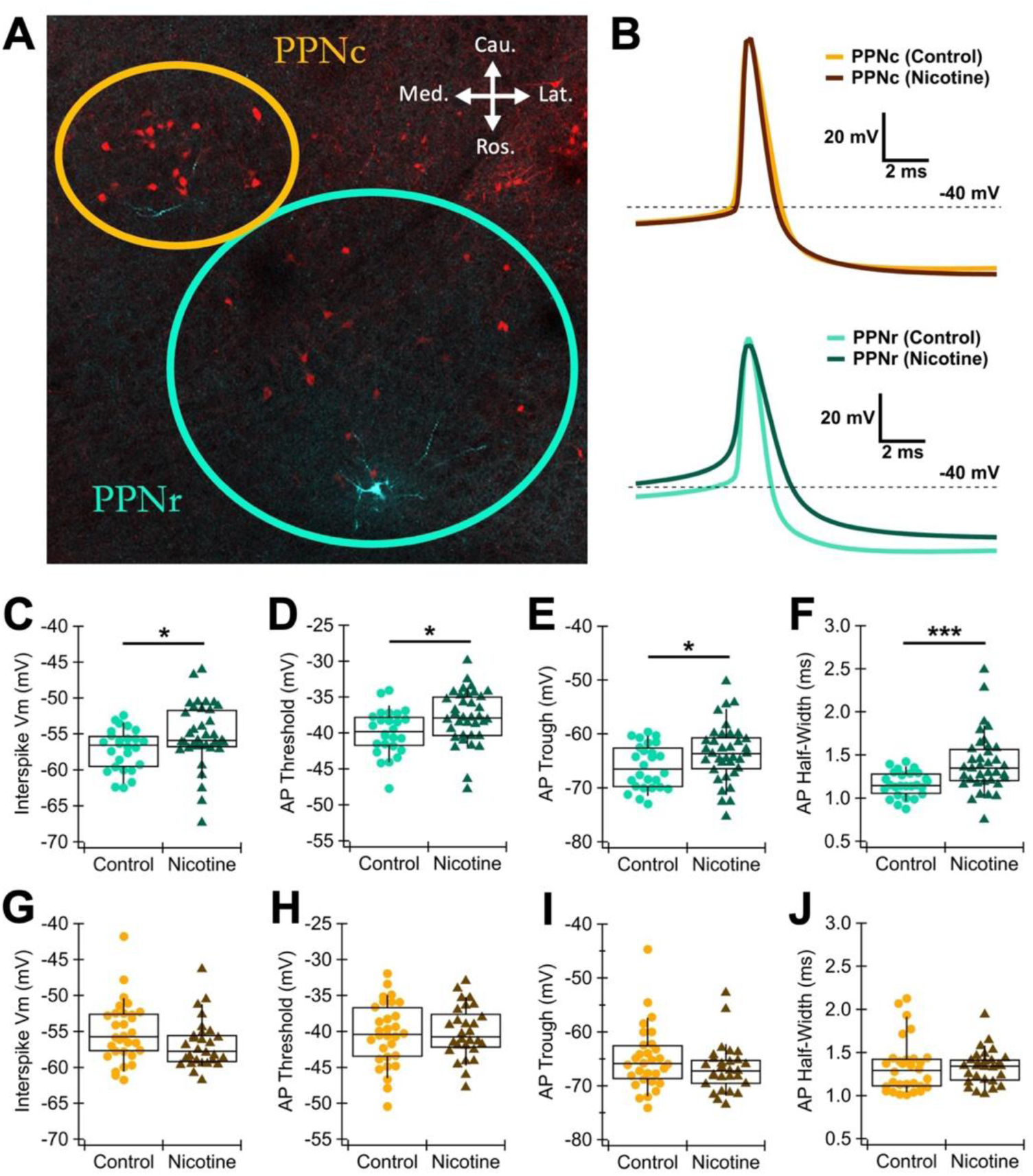
PPN cholinergic (ACh) neurons exhibit subregion-specific changes in action potential (AP) waveform after chronic nicotine. **(A)** Confocal overview image of the PPN on a horizontal slice, showing ChAT+ neurons (red, ChAT-TdTomato) and patched neurons filled with neurobiotin (cyan, streptavidin-Cy5); the yellow circle indicates the approximate boundary of the caudal PPN (PPNc) whereas the blue-green circle, the rostral PPN (PPNr). Cau = caudal, Med = medial, Ros = rostral, and Lat = lateral. **(B)** Representative AP waveforms of PPN ACh neurons averaged from spontaneous pacemaking APs over 30 s, color-coded by subregion (rostral: blue-green tones; caudal: yellow-brown tones) and treatment (control: light; nicotine: dark). **(C)** Summary of PPNr (control: light blue-green; nicotine: dark green) interspike Vm, **(D)** AP threshold, **(E)** hyperpolarization trough, and **(F)** half-width. **(G)** Summary of PPNc (control: yellow; nicotine: brown) interspike Vm, **(H)** AP threshold, **(I)** hyperpolarization trough, and **(J)** half-width.

However, chronic nicotine significantly altered AP shape in the rostral but not caudal PPN (Figure 5B). In rostral PPN ACh neurons, chronic nicotine significantly depolarized the interspike membrane potential (Vm) (control: –57.17 ± 0.56, N = 9, n = 26; nicotine: –55.20 ± 0.73, N = 13, n = 36; t = -2.15, p = 0.0360, t-test; Figure 5C), AP threshold (control: –39.92 ± 0.61; nicotine: –37.84 ± 0.61; t = -2.41, p = 0.0193, t-test; Figure 5D), and AP trough (control: –66.10 ± 0.81; nicotine: –63.61 ± 0.89; t = -2.07, p = 0.0427, t-test; Figure 5E). Nicotine also significantly increased AP half-width (control: 1.16 ± 0.03; nicotine: 1.40 ± 0.06; W = 230, p = 0.0005, Wilcoxon test; Figure 5F). By contrast, the caudal PPN showed no significant differences in interspike Vm (control N = 10, n = 30; nicotine N = 13, n = 28; W = 306, p = 0.0772; Wilcoxon test; Figure 5G), threshold (t = -0.20, p = 0.8420, t-test; Figure 5H), trough (W = 332, p = 0.1747, Wilcoxon test; Figure 5I), and half-width (W = 385, p = 0.5940, Wilcoxon test; Figure 5J) between control and nicotine groups, although AP peak amplitude was modestly increased (t = -2.10, p = 0.0409; t-test; Figure S4).

These results indicate that chronic nicotine selectively depolarizes and broadens action potentials in rostral PPN ACh neurons without altering firing rates or passive membrane properties. AP width broadening and interspike voltage depolarization are generally associated with enhanced Ca^2+^ influx and neurotransmitter release (Hoppa et al., 2014; Begum et al., 2016), suggesting that chronic nicotine might increase PPN ACh output to downstream targets such as SNc or striatum.

### Chronic nicotine selectively decreases spike frequency adaptation in caudal PPN cholinergic neurons

Two of the defining electrophysiological features that distinguish ACh neurons from non-ACh neurons in the PPN are strong spike frequency adaptation (SFA) and a prominent M-current (Bordas et al., 2015; Petzold et al., 2015). Both properties are thought to regulate afterhyperpolarization trough and phasic firing patterns critical for network synchrony and sensory gating (Bordas et al., 2015; Petzold et al., 2015). We therefore examined whether SFA and M-current amplitude in PPN ACh neurons are altered by chronic nicotine.

As with AP shape (Figure 5), nicotine differentially altered these properties in the caudal vs rostral PPN. Using a series of depolarizing current steps (+50 to +200 pA, 1 s), we found that nicotine-treated caudal PPN neurons exhibited reduced SFA compared to controls across all current steps (Figure 6A), reaching statistical significance at 150 (control: N = 10, n = 32; nicotine: N = 13, n = 29; t = 2.29, p = 0.0259; t-test) and 200 pA steps (t = 2.47, p = 0.0162; t-test; Figure 6B,C). In contrast, rostral PPN neurons showed no significant differences in adaptation index between groups (Figure 6D,E). Thus, chronic nicotine selectively reduces SFA in caudal PPN ACh neurons which would increase sustained firing during prolonged excitation and decrease burst propensity.

**Figure 6.**
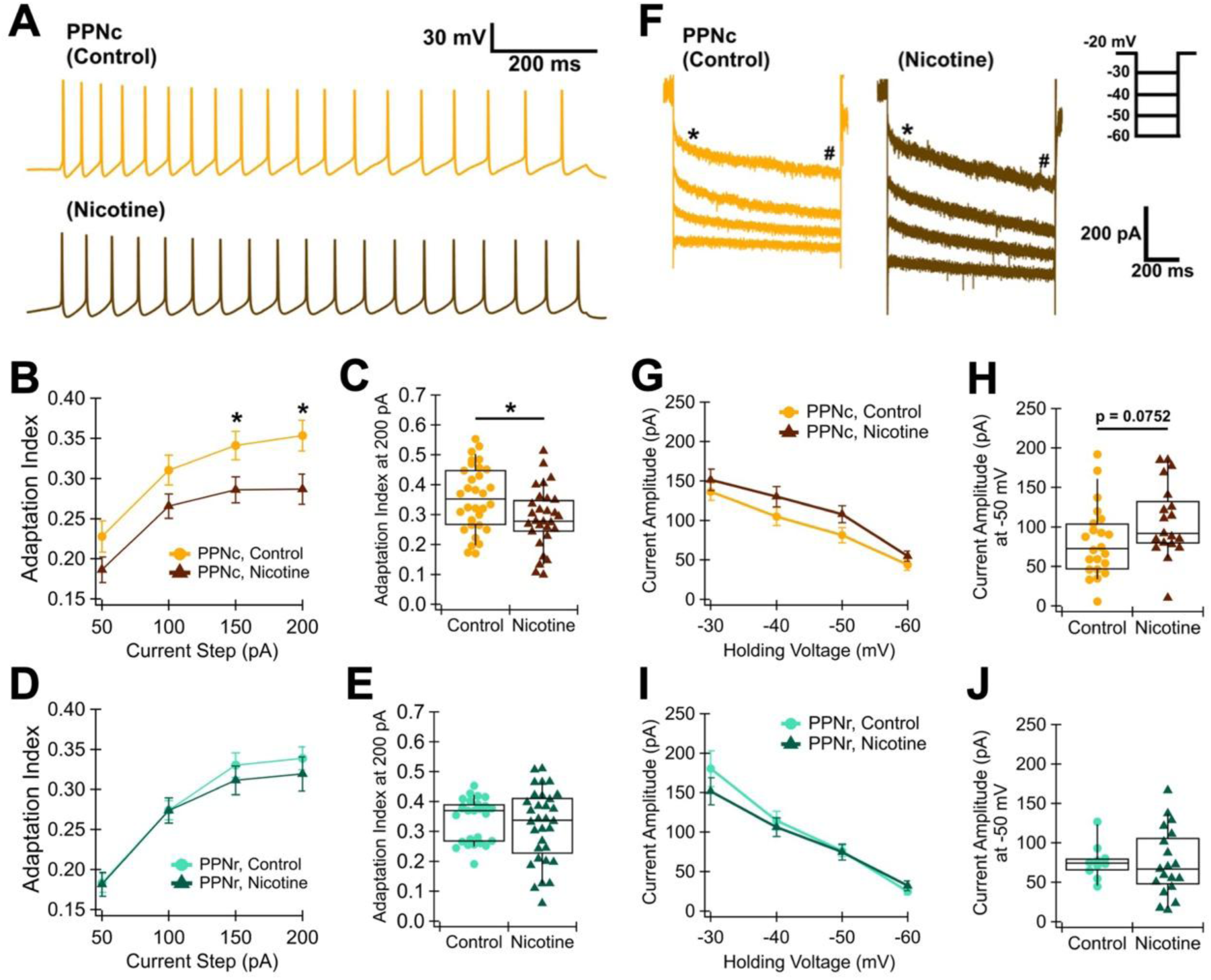
PPN ACh neurons exhibit subregion-specific reduction of SFA after chronic nicotine. **(A)** Representative caudal PPN (PPNc) action potential trains elicited by a 100-pA current step, illustrating SFA as progressive slowing of firing rates during sustained depolarization, from control (yellow) and nicotine (brown) groups. **(B)** Averaged adaptation index of PPNc neurons elicited by depolarizing current steps. **(C)** Summary of PPNc neuron adaptation index at the 200-pA step. **(D)** Averaged adaptation index of rostral PPN (PPNr) neurons elicited by depolarizing current steps. **(E)** Summary of PPNr neuron adaptation index at the 200-pA step. **(F)** Representative PPNc M-current traces elicited by 1-s repolarizing steps from –20 mV holding potential to –30, –40, –50, and –60 mV; the asterisk (*) indicates the early steady state current and the hash (#) indicates the late steady state current, with M-current amplitude calculated as the difference between the two. **(G)** Averaged PPNc M-current amplitude elicited by repolarizing steps from –20 mV. **(H)** Summary of PPNc M-current amplitude at the –50 mV step. **(I)** Averaged PPNr M-current amplitude elicited by repolarizing steps from –20 mV. **(J)** Summary of PPNr M-current amplitude at the –50 mV step.

M-currents are slow, non-inactivating subthreshold K^+^ conductances mediated by Kv7 (KCNQ) channels (Gu et al., 2005). Consistent with their classical functions in regulating excitability, prior studies have shown that M-currents in PPN ACh neurons deepen the afterhyperpolarization trough and enhance SFA (Bordas et al., 2015; Bayasgalan et al., 2021). To test whether changes in M-currents underlie the nicotine-induced reduction in adaptation, we measured M-current deactivation using voltage steps (–30 to –60 mV, 1 s) from a holding potential of –20 mV (Figure 6F). The average M-current amplitude did not differ significantly between control and nicotine-treated neurons in either subregion (Figure 6G), but caudal PPN neurons from nicotine-treated mice showed a trend toward larger M-currents, most prominent at –50 mV (control: N = 10, n = 22; nicotine: N = 12, n = 19; t = -1.83, p = 0.0752, t-test; Figure 6H). In the rostral PPN, the M-current amplitude did not differ between groups (Figure 6I,J).

This result was unexpected, as enhanced M-currents would typically promote rather than reduce SFA in PPN ACh neurons (Bordas et al., 2015; Bayasgalan et al., 2021). The discrepancy suggests that the diminished adaptation is unlikely to result from reduced M-current activity, and may instead reflect compensatory changes secondary to M-current upregulation or involvement of other ionic mechanisms. To investigate this, we correlated adaptation index with M-current amplitude across individual neurons. Larger M-currents were associated with stronger adaptation in both control and nicotine conditions, confirming their expected functional relationship (Figure S5). Therefore, chronic nicotine appears to reduce SFA through mechanisms independent of M-current modulation – potentially through altered calcium-dependent potassium conductances or Na^+^ channel inactivation, which are also involved in the regulation of SFA (Vandecasteele et al., 2011; Vandael et al., 2012; Upchurch et al., 2022).

### Chronic nicotine reduces dendritic length in rostral PPN and accelerates kinetics of excitatory synaptic inputs to caudal PPN

Chronic nicotine alters dendritic branching in striatal neurons (McDonald et al., 2005; Ehlinger et al., 2016, 2017). We therefore hypothesized that chronic nicotine might induce structural and synaptic remodeling in PPN ACh neurons in addition to altering intrinsic properties. Dendritic reconstruction and excitatory synaptic input analyses revealed that chronic nicotine differentially affected the rostral and caudal PPN subregions (Figure 7A–H). Quantification of the area under the curve (AUC) of the Sholl profiles showed a significant reduction in the rostral PPN after chronic nicotine exposure, indicative of dendritic pruning (control: 187.61 ± 7.25 intersections·µm, N = 10, n = 27; nicotine: 161.04 ± 8.84 intersections·µm, N = 12, n = 28; t = 2.33, p = 0.0240, t-test; Figure 7G). Total dendritic length in the rostral PPN neurons also trended lower in nicotine-treated mice (control: 2210 ± 164 µm, nicotine: 2075 ± 119 µm, t = 1.98, p = 0.0527, t-test; Figure 7H). By contrast, caudal PPN neurons showed no difference in either Sholl AUC (W = 202.5, p = 0.8515, Wilcoxon test; Figure 7D) or total dendritic length (W = 206, p = 0.9281, Wilcoxon test; Figure 7E) between groups. These findings show that nicotine selectively reduces dendritic complexity in rostral PPN ACh neurons.

**Figure 7.**
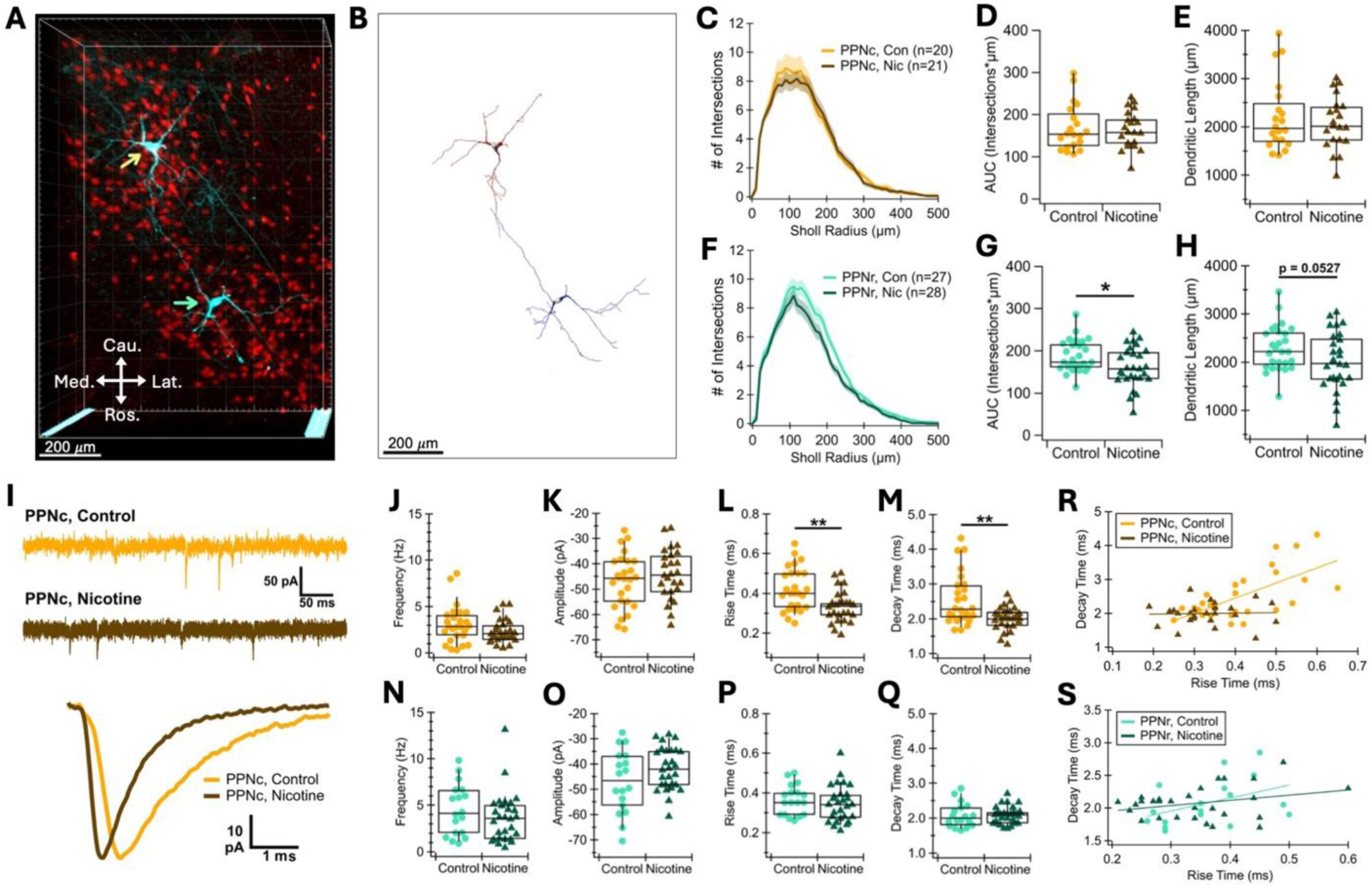
PPN ACh neurons exhibit subregion-specific dendritic pruning and faster sEPSC gating kinetics after chronic nicotine. **(A)** Three-dimensional confocal Z-stack image shown in Imaris of two patched PPN ACh neurons (cyan, neurobiotin-filled, streptavidin-Cy5 stain) against a background of neighboring ChAT^+^ neurons (red, ChAT-TdTomato), with yellow and blue-green arrows indicating somas of caudal PPN (PPNc) and rostral PPN (PPNr) neurons, respectively, on a horizontal brain slice. **(B)** The same two PPN ACh neurons shown in (A) with soma and dendrites digitally reconstructed in Imaris. **(C)** Sholl analysis of PPNc (control: yellow; nicotine: brown) with the number of dendritic intersections plotted against Sholl radius value of 10 μm. **(D)** Area under the curve (AUC) of PPNc Sholl graph shown in C. **(E)** Summary of total dendritic length of PPNc. **(F)** Sholl analysis of PPNr (control: blue-green, nicotine: dark green) neurons with the number of dendritic intersections plotted against Sholl radius value of 10 μm. **(G)** Area under the curve (AUC) of PPNr Sholl graph shown in F. **(H)** Summary of total dendritic length of PPNr. **(I)** Representative voltage-clamp recording of sEPSCs at –70 mV holding potential (top) and averaged sEPSC waveform (bottom) from one control (yellow) and one nicotine (brown) PPNc neuron. **(J)** Summary of PPNc sEPSC frequency, **(K)** amplitude, **(L)** rise time, and **(M)** decay time. **(N)** Summary of PPNr frequency, **(O)** amplitude, **(P)** rise time, and **(Q)** decay time. **(R)** Decay times plotted against rise times in individual PPNc and **(S)** PPNr neurons. Data from each subregion and treatment group were fitted with linear regression lines.

Excitatory synaptic transmission was assessed by recording sEPSCs in voltage-clamp configuration holding at -70 mV. PPN ACh neurons exhibited frequent synaptic currents with moderate amplitudes, suggesting a strong overall excitatory drive onto these cells (Figure 7I). In the caudal PPN, the average sEPSC frequency and amplitude were not significantly altered by chronic nicotine (control: N = 10, n = 26, nicotine: N = 13, n = 26; frequency, W = 256.5, p = 0.1379, Wilcoxon test; amplitude, t = -0.89, p = 0.3797; t-test; Figure 7J,K), but both rise and decay times were significantly faster in neurons from nicotine-treated mice (rise time: control 0.42 ± 0.02 ms, nicotine 0.33 ± 0.02 ms, t = 3.40, t-test; decay time: control 2.54 ± 0.14 ms, p = 0.0014, nicotine 2.00 ± 0.06 ms, W = 185.5, p = 0.0046, Wilcoxon test; Figure 7L,M). In contrast, rostral PPN showed no significant changes in any sEPSC parameter (frequency, W = 190.5, p = 0.1697, Wilcoxon test; amplitude, t = -1.50, p = 0.1453; t-test; rise time, t = 0.77, p = 0.4444, t-test; decay time, t = 0.09, p = 0.9325, t-test: Figure 7N–Q).

sEPSC rise time and decay time are biophysical properties that normally co-vary due to synaptic location and dendritic filtering properties (McBain and Dingledine, 1992; Evans et al., 2012). To determine whether nicotine alters the relationship between these properties, we performed within-cell correlations between rise and decay times. Results revealed that the control caudal PPN ACh neurons showed a strong positive relationship (r = 0.64, p = 0.0003), whereas this relationship was flattened following chronic nicotine (r = 0.04, p = 0.8584; comparison of control vs nicotine slopes, p = 0.0136,

Fisher’s r-to-z test; Figure 7R). In rostral PPN, the rise-decay correlation was modest and trending in controls (r = 0.43, p = 0.0737), and nicotine did not significantly affect the slope (r = 0.29, p = 0.1301; r of control vs nicotine, p = 0.6244, Fisher’s r-to-z transformation test; Figure 7S). These results show that chronic nicotine induces subtle yet significant region-specific plasticity within the PPN: it reduces dendritic arborization in rostral PPN while accelerating excitatory synaptic kinetics in the caudal PPN. The faster sEPSC kinetics in the caudal PPN, together with the loss of correlation between rise and decay times, suggests that chronic nicotine alters postsynaptic receptor characteristics rather than passive electronic filtering. The absence of nicotine-induced changes in whole-cell capacitance (Figure S4K) and total dendritic length (Figure 7E) in the caudal PPN further supports this interpretation.

## Discussion

Here we show that chronic nicotine exposure reshapes the physiology of two brainstem neuronal populations central to PD pathology. In SNc DA neurons, nicotine suppresses excitability by lowering spontaneous pacemaking frequency, diminishing burst and rebound firing propensity, and decoupling tonic firing rate from proximal dendritic calcium influx. In contrast, the PPN underwent complex subregion-specific remodeling: the rostral ACh neurons exhibited membrane potential depolarization, action potential broadening, and dendritic pruning, whereas the caudal ACh neurons showed reduced SFA and accelerated sEPSC kinetics. This is the first study to demonstrate that chronic nicotine drives intrinsic, synaptic, and morphological plasticity in adult PPN ACh neurons, uncovering pronounced rostro-caudal divergence. These adaptations in SNc excitability and PPN remodeling are potential cellular mechanisms for the well-established inverse correlation between chronic nicotine and PD risk in epidemiology and preclinical animal models.

We found that chronic nicotine consistently reduced spontaneous tonic firing frequency in SNc DA neurons from adult male and female mice, extending prior observations from a shorter 10-day exposure paradigm in young mice of unspecified sex (Nashmi et al., 2007; Xiao et al., 2009). Those studies attributed the firing reduction to enhanced GABAergic inhibition from the substantia nigra pars reticulata. Since we did not observe changes in excitatory synaptic drive, an increased inhibitory drive may also contribute here. However, we found diminished burst propensity and rebound firing frequency, suggesting that there is also nicotine-induced plasticity of intrinsic characteristics. The trending reduction in hyperpolarization-induced afterdepolarization (ADP) suggests that T-type calcium channel activity may be downregulated by chronic nicotine. T-type calcium channels generate the ADP, contribute to rebound bursts, and their strong expression is a defining characteristic of vulnerable SNc neurons (Evans et al., 2017). Although ADP sampling is sensitive to slice orientation, the consistent rebound and burst suppression across neurons aligns with a protective dampening of T-type-mediated excitability in response to chronic nicotine.

We had originally hypothesized that chronic nicotine would reduce SNc dendritic L-type calcium signals, but found that L-type channel contribution to dendritic calcium influx was not changed significantly after nicotine exposure. Importantly, while the overall somatodendritic calcium signal was not altered by nicotine, the fluorescence method used here only allows for comparison of relative and not absolute calcium concentration. However, it does allow for the evaluation of the nifedipine-sensitive L-type-mediated components and the relationship between firing rate and dendritic calcium influx. This relationship that is supralinear in SNc neurons (Hage and Khaliq, 2015), and our finding that nicotine flattens this relationship suggest that nicotine buffers the subpopulation of SNc neurons with higher firing rates against excessive dendritic calcium loading, potentially mitigating mitochondrial oxidative stress implicated in selective vulnerability (Guzman et al., 2010, 2018; Dryanovski et al., 2013). However, it also suggests that low firing rates cause higher dendritic calcium loads after chronic nicotine. The robust nifedipine effects in SNc neurons of both control and nicotine-treated mice are consistent with those we have reported previously (Chen and Evans, 2024) and point to the possibility that nicotine alters non-L-type channel activity in SNc DA neurons.

Although chronic nicotine increases dendritic arborization in striatal spiny projection neurons of the dorsolateral striatum (Ehlinger et al., 2017) and the nucleus accumbens shell (McDonald et al., 2005; Ehlinger et al., 2016), our findings reveal a notable absence of such effects in SNc DA neurons. The lack of remodeling persisted when neurons were categorized by electrophysiological subtypes, with no differences in arbor complexity between ADP^+^ (vulnerable ventral-tier) and ADP^-^ (resilient) subpopulations. Prior work on nicotine’s neuroprotective actions in PD models has emphasized dopaminergic soma counts, biochemical markers, or terminal release following toxin-induced lesion (Bordia et al., 2006; Quik et al., 2006; Khwaja et al., 2007; Huang et al., 2009), but morphological assessments were lacking. Our results suggest that chronic nicotine affects SNc DA neurons primarily through functional adaptations (i.e., dampening excitability and calcium load), rather than structural remodeling as is observed in striatal neurons.

In the PPN, chronic nicotine induced differential plasticity in the rostral and caudal subregions. This supports the emerging understanding of regional PPN differences in function and connectivity (Dautan et al., 2014; Baksa et al., 2019; Huerta-Ocampo et al., 2020, 2021; Fallah et al., 2025; Sharma et al., 2025). The rostral PPN preferentially innervates motor-related basal ganglia circuits including the SNc and dorsolateral striatum (Martinez-Gonzalez et al., 2011; Mena-Segovia and Bolam, 2017; Xiao et al., 2020), making nicotine’s effects here particularly relevant to PD pathology. We observed selective alterations in rostral PPN ACh neurons: depolarized AP threshold and interspike Vm, increased AP half-width, and reduced dendritic complexity. AP broadening may enhance Ca^2+^ influx and acetylcholine release at terminals (Hoppa et al., 2014; Begum et al., 2016; Kramer et al., 2020). Enhanced rostral cholinergic drive might promote SNc DA neuron survival through reciprocal connectivity (McGeer and McGeer, 1984; Bensaid et al., 2016; Sharma et al., 2020) or muscarinic receptor-mediated reduction of rebound-induced Ca^2+^ influx (Beaver and Evans, 2025) and AMPA receptor currents (Campos et al., 2025). Alternatively, it could ameliorate PD symptoms linked to cholinergic loss, such as gait deficits, sleep disturbances, and cognitive impairments (Bohnen et al., 2009; Perez-Lloret and Barrantes, 2016; Chambers et al., 2019). However, somatic AP changes may not faithfully propagate to terminals, tempering these interpretations (Hu and Bean, 2018; Kramer et al., 2020; Gonzalez Sabater et al., 2021).

In the caudal PPN, chronic nicotine accelerated sEPSC rise and decay times selectively in caudal PPN ACh neurons, without altering frequency and amplitude. Fast glutamatergic transmission in central neurons is predominantly mediated by AMPA receptors (AMPAR), and the subunit composition of these receptors affects rise and decay times (Geiger et al., 1997; Isaac et al., 2007). GluA2-lacking (Ca^2+^-permeable) receptors exhibit faster channel deactivation kinetics than GluA2-containing counterparts (Geiger et al., 1995). Cerebellar stellate cells express GluA2-lacking AMPARs that that shift to GluA2-containing subtypes after fear conditioning, shifting the median decay time constants of sEPSC from 0.5 to 0.7 ms without altering current amplitude (Savtchouk and Liu, 2011). This corresponds to a ∼29% change in decay kinetics, comparable to the ∼21% reduction in decay we observed in caudal PPN ACh neurons. Importantly, nicotinic receptor activation has been linked to upregulation of GluA2-lacking AMPARs (or downregulation of GluA2-containing AMPARs) in the VTA (Vieyra-Reyes et al., 2008; Gao et al., 2010), PFC (Tang et al., 2015), and NAc synaptosomes (Marchi et al., 2015). A similar mechanism could underlie the acceleration in sEPSC kinetics observed here. On the other hand, TARP auxiliary subunit subtypes also tune AMPAR gating kinetics, with γ-2 coupling to faster currents and γ-4/8 producing slower currents (Milstein et al., 2007). Future experiments will be needed to determine the mechanism underlying the nicotine-induced increase in EPSC gating kinetics. Together with reduced SFA, these chronic nicotine-mediated alterations could sustain efficient cholinergic output from the caudal PPN.

In addition to its established role in motor control and Parkinsonian vulnerability, the PPN provides cholinergic modulation that enhances signal-to-noise ratios in reward processing and sensory gating (Picciotto et al., 2012). The caudal PPN, with projections to both motor and limbic/associative structures (Mena-Segovia and Bolam, 2017), makes nicotine-induced plasticity in this subregion likely to influence reward and attentional circuits. Computational models suggest that SFA promotes metabolic efficiency at the cost of history-dependent coding in sensory networks (Gutierrez and Denève, 2019). By reducing SFA and accelerating sEPSC kinetics selectively in caudal PPN ACh neurons, chronic nicotine may shift this tradeoff: increasing metabolic demands while enabling faster, more regular firing and amplified responses to sustained inputs. At the systems level, these adaptations could heighten vigilance in sensory-attentional tasks (Pan and Hyland, 2005; Hong and Hikosaka, 2014), disrupt sharp sleep-to-wake transitions (Petzold et al., 2015), alter nicotine reward cue processing (Laviolette et al., 2002), and influence PD progression by shifting energetic burden or alpha/beta oscillatory synchrony (Thevathasan et al., 2012; Li and Zhang, 2015). Thus, the net effects of nicotine-induced PPN changes likely depend on temporally and behaviorally sensitive activation of this cell population. Ultimately, future investigation is needed to determine whether chronic nicotine’s subregion-specific remodeling of PPN physiology is adaptive or maladaptive across different behavioral and pathological context.

Together, this study illuminates cellular adaptations that contribute to nicotine’s complex influence on the brainstem neurons vulnerable to neurodegeneration and implicated in psychiatric disorders. The extensive chronic nicotine administration paradigm and the use of adult male and female mice reveal novel mechanisms by which nicotine can influence sensorimotor circuitry in the extended basal ganglia. It is important to note that our recordings were performed in acute brain slice, preserving intrinsic properties and local synaptic inputs but excluding network influences. Future work using *in vivo* methods will be critical for understanding the time course of these effects and how they contribute to neural processing in the intact brain. Importantly, additional experiments building on this study should determine whether the intrinsic, synaptic, and morphological changes caused by chronic nicotine are protective or harmful to circuit function.

## Acknowledgements

This work was supported by the American Parkinson’s Disease Association Research Grant 2021APDA00RG00000209666, Parkinson’s Foundation Stanley Fahn Junior Faculty Award #PF-SF-JFA-1040267, and BRAIN Initiative K99/R00 award #R00NS112417 awarded to RCE. After the original draft was written by Dr. Chen, LLM AI tools were used for language editing. Final manuscript was edited by Drs. Evans and Chen. We thank members of the Evans Lab for feedback on earlier versions of this manuscript.

