## Supplemental Figures for "Chronic nicotine reduces nigral dopaminergic activity and remodels pedunculopontine cholinergic subpopulations"

Supplementary Figures

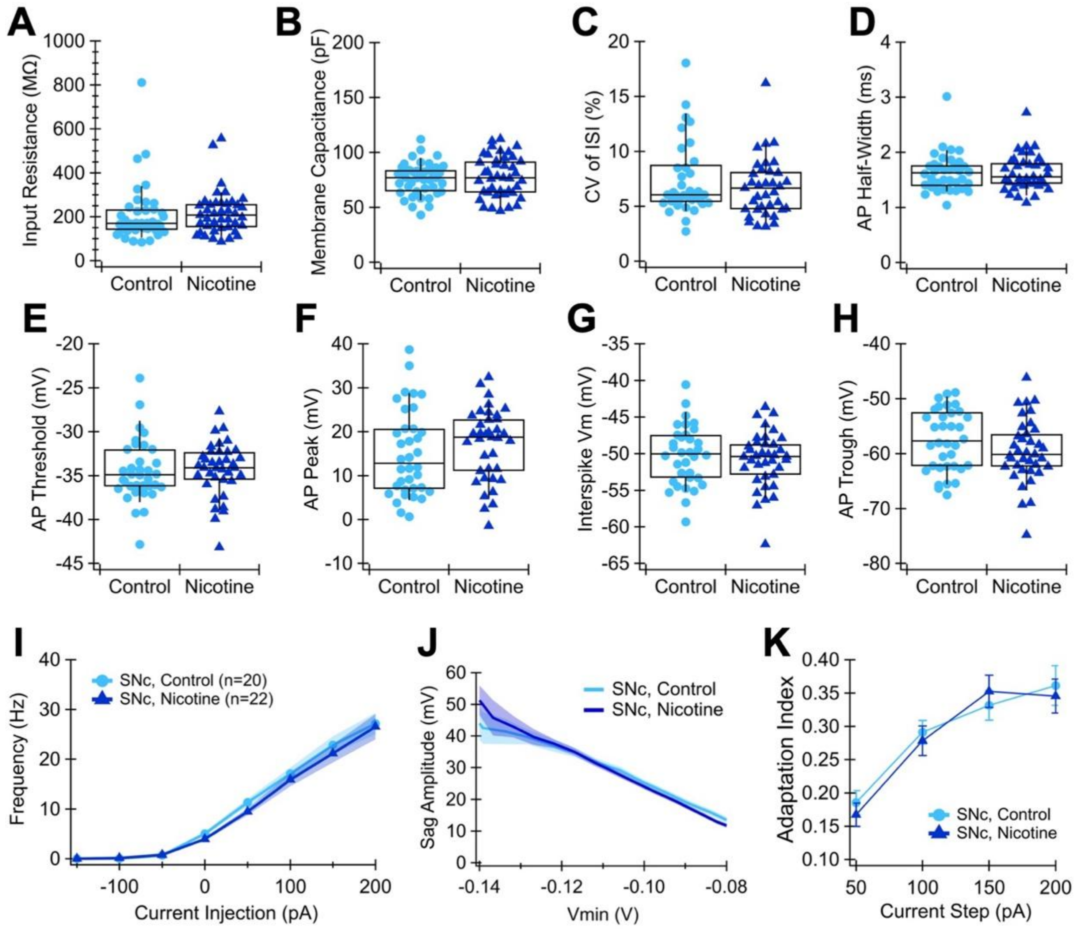


**Figure S1.** SNc dopaminergic (DA) neurons show no changes in most firing properties after chronic nicotine. **(A)** Input resistance (control n = 42, nicotine n = 41, p = 0.2740, Wilcoxon test) and **(B)** whole-cell capacitance (p = 0.6000, t-test), measured by a voltage step from –70 to –75 mV (100 ms duration) applied to SNc DA neurons in control (light blue) and nicotine (dark blue) groups. **(C)** Firing regularity (control n = 36, nicotine n = 37, p = 0.4389, Wilcoxon test), calculated as coefficient of variation of the interspike interval (CoV of ISI), **(D)** AP half-width (p = 0.8392, Wilcoxon test), **(E)** threshold (p = 0.4455, Wilcoxon test), **(F)** peak (p = 0.3449, t-test), **(G)** interspike Vm (p = 0.5000, Wilcoxon test), and **(H)** trough (p = 0.1445, Wilcoxon test) of SNc DA neurons averaged from spontaneous pacemaking activity over 30 s. **(I)** Current-frequency (I-f) relationship of SNc DA neurons measured by current steps (–150 to +200 pA, 50 pA increments, 1 s duration). **(J)** Hyperpolarization-induced voltage sag of SNc DA neurons elicited by current steps (ranging from –300 to –700 pA, 100 pA increments, 1 s duration), calculated by subtracting the most hyperpolarized minimal voltage (Vmin) from the steady state potential reached by the end of each current step, plotted against Vmin as an indicator of HCN channel activity. **(K)** Spike frequency adaptation of SNc DA neurons, calculated as the average adaptation index elicited by the depolarizing current steps described in **I**.

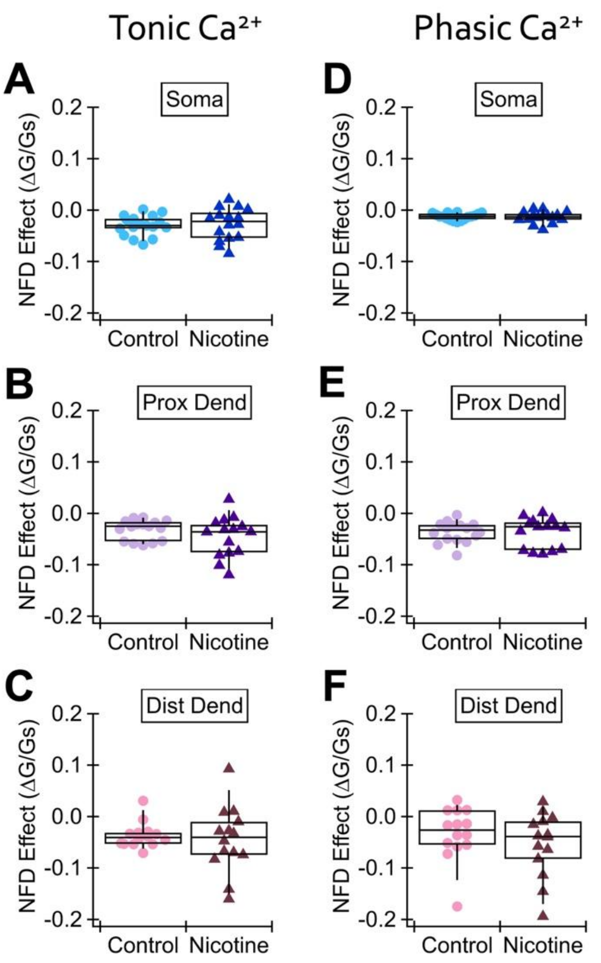


**Figure S2.** SNc DA neurons show no changes in L-type calcium channel-mediated dendritic calcium signals after chronic nicotine. **(A)** Difference between tonic $Ca^{2+}$ signals before and after bath treatment with the L-type calcium channel blocker nifedipine (10 μM), measured at the soma (control n = 17, nicotine n = 16, p = 0.8400, t-test), **(B)** proximal dendrite (control n = 16, nicotine n = 15, p = 0.2020, Wilcoxon test), and **(C)** distal dendrite (control n = 14, nicotine n = 14, p = 0.8388, Wilcoxon test) of SNc DA neurons. **(D)** Difference between phasic $Ca^{2+}$ signals before and after nifedipine, measured at the soma (p = 0.6495, t-test), **(E)** proximal dendrite (p = 0.8001, Wilcoxon test), and **(F)** distal dendrite (p = 0.3287, Wilcoxon test) of SNc DA neurons.

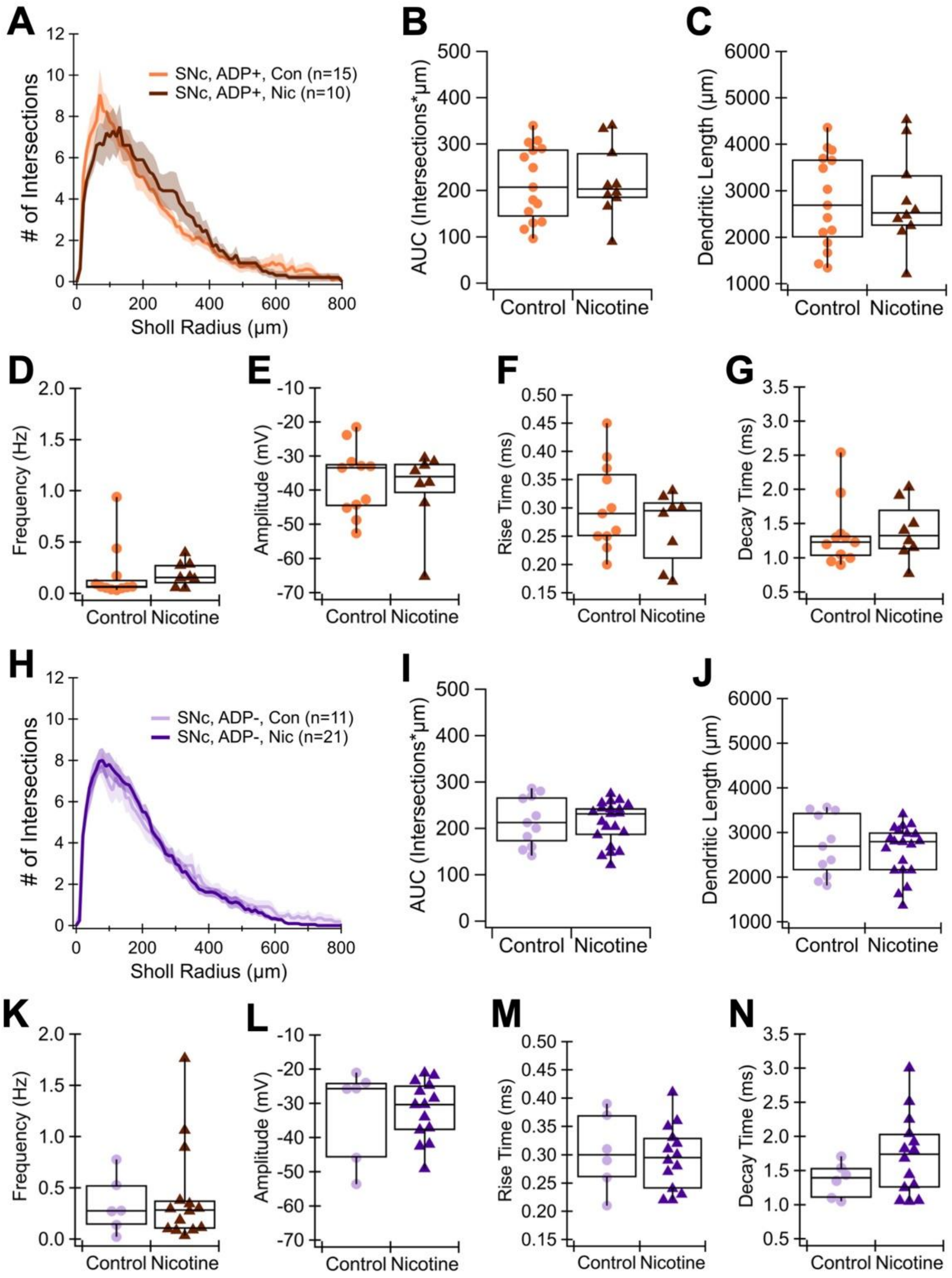


**Figure S3.** Chronic nicotine does not change the dendritic morphology and excitatory synaptic drive of $ADP^+$ and $ADP^-$ SNc DA neurons. **(A)** Sholl analysis of $ADP^+$ SNc DA neurons, with the number of dendritic intersections plotted against Sholl radius value of 10 μm, in control (orange) and nicotine (brown) groups. **(B)** Area under the curve (AUC) of $ADP^+$ Sholl graph (control n = 15, nicotine n = 10, p = 0.8977, t-test) and **(C)** total dendritic length shown in **A** (p = 0.8005, t-test). **(D)** $ADP^+$ sEPSC frequency (control n = 11, nicotine n = 8, p = 0.2997, Wilcoxon test), **(E)** amplitude (p = 0.9678, Wilcoxon test), **(F)** rise time (p = 0.2595, t-test), and **(G)** decay time (p = 0.5854, Wilcoxon test). **(H)** Sholl analysis of $ADP^-$ SNc DA neurons, with the number of dendritic intersections plotted against Sholl radius value of 10 μm, in control (light purple) and nicotine (dark purple) groups. **(I)** Area under the curve (AUC) of ADP$^-$ Sholl graph (control n = 11, nicotine n = 21, p = 0.9741, t-test) and **(J)** total dendritic length shown in **H** (p = 0.6246, t-test). **(K)** ADP$^-$ sEPSC frequency (control n = 6, nicotine n = 14, p = 0.9187, Wilcoxon test), **(L)** amplitude (p = 0.9318, t-test), **(M)** rise time (p = 0.7584, t-test), and **(N)** decay time (p = 0.071, t-test).

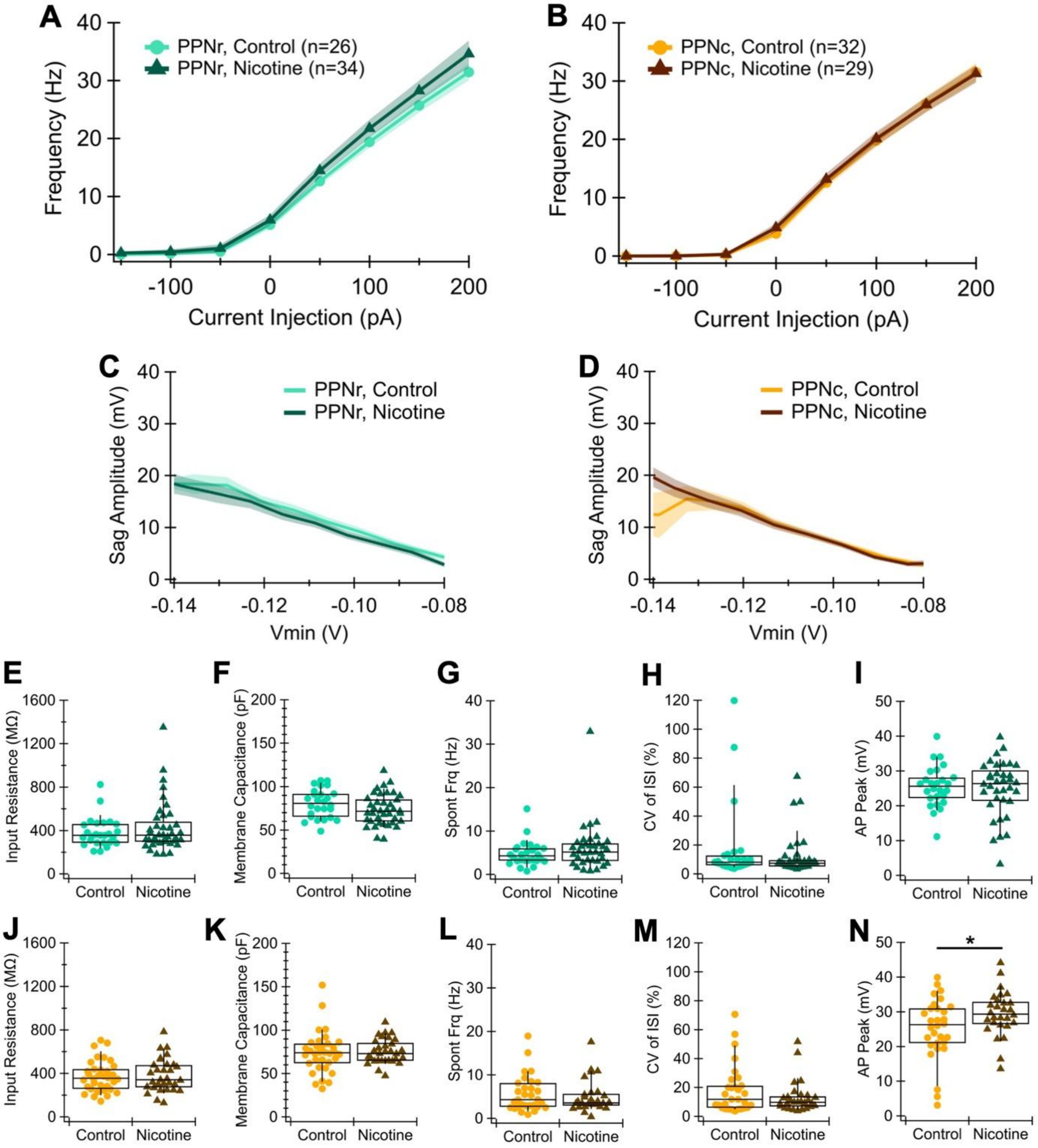


**Figure S4.** PPN cholinergic (ACh) neurons show no changes in current-frequency relationship, HCN channel activity, pacemaking frequency, and other passive properties, except AP peak in caudal PPN showing modest increase. **(A)** Current-frequency (I-f) relationship of rostral PPN (PPNr) and **(B)** caudal PPN (PPNc) ACh neurons measured by current steps (–150 to +200 pA, 50 pA increments, 1 s duration). **(C)** Hyperpolarization-induced voltage sag elicited by current steps (ranging from –300 to –700 pA, 100 pA increments, 1 s duration), calculated by subtracting the most hyperpolarized minimal voltage (Vmin) from the steady state potential reached by the end of each current step, plotted against Vmin as an indicator of HCN channel activity in PPNr and

**(D)** PPNc neurons. **(E)** Input resistance (control n = 26, nicotine n = 39, p = 0.8785, Wilcoxon test) and **(F)** whole-cell capacitance (p = 0.0835, t-test), measured by a voltage step from –70 to –75 mV (100 ms duration) applied to PPNr neurons in control (light blue-green) and nicotine (dark blue-green) groups. **(G)** Spontaneous pacemaking frequency (control n = 26, nicotine n = 36, p = 0.2470, Wilcoxon test), **(H)** firing regularity, calculated as coefficient of variation of the interspike interval (CoV of ISI) (p = 0.2902, Wilcoxon test), and **(I)** AP peak of PPNr neurons averaged from spontaneous pacemaking activity over 30 s (p = 0.6958, t-test). **(J)** Input resistance (control n = 34, nicotine n = 31, p = 0.8198, t-test) and **(K)** whole-cell capacitance of PPNc neurons in control (yellow) and nicotine (brown) groups (p = 0.7296, Wilcoxon test). **(L)** Spontaneous pacemaking frequency (control n = 30, nicotine n = 28, p = 0.6486, t-test), **(M)** firing regularity, calculated as coefficient of variation of the interspike interval (CoV of ISI) (p = 0.4080, Wilcoxon test), and **(N)** AP peak of PPNc neurons averaged from spontaneous pacemaking activity over 30 s (p = 0.0409, t-test).

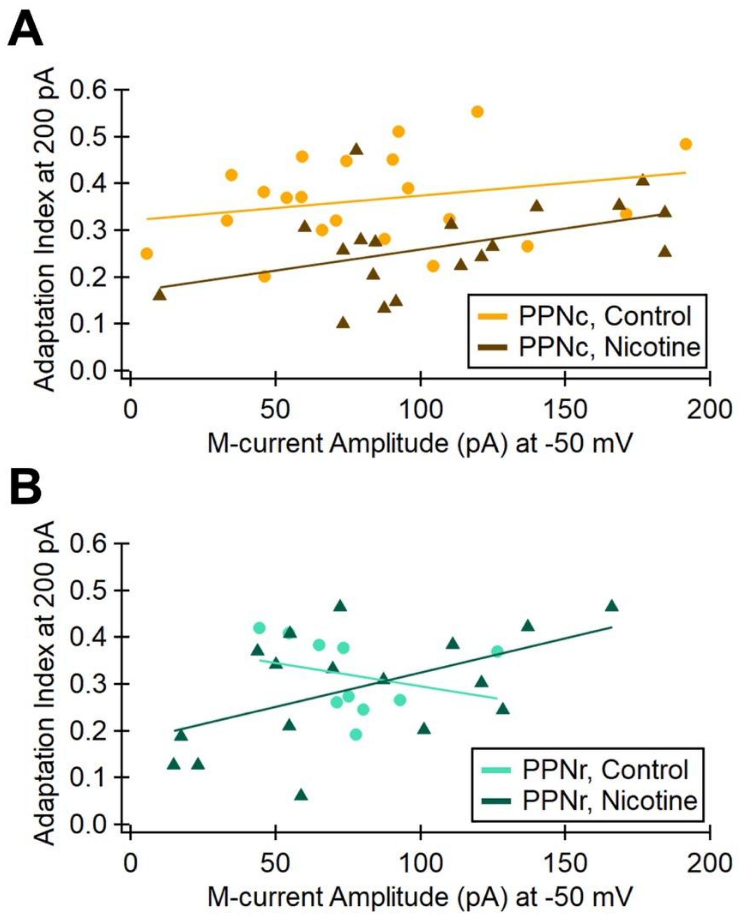


**Figure S5.** M-current amplitude and adaptation index are positively correlated in PPN ACh neurons. **(A)** Caudal PPN (PPNc) adaptation index at the 200-pA step plotted against the M-current amplitude at the –50-mV step in control (yellow) and nicotine (brown) groups (control n = 21, r = 0.2515, p = 0.2714; nicotine n = 19, r = 0.4426, p = 0.0577). **(B)** Rostral PPN (PPNr) adaptation index at the 200-pA step plotted against the M-current amplitude at the –50-mV step in control (light blue-green) and nicotine (dark blue-green) groups (control n = 10, r = -0.2775, p = 0.4378; nicotine n = 17, r = 0.5264, p = 0.0300).
